# Novel tri-segmented rhabdoviruses: A data mining expedition unveils the cryptic diversity of cytorhabdoviruses

**DOI:** 10.1101/2023.11.06.565797

**Authors:** Nicolás Bejerman, Ralf Georg Dietzgen, Humberto Debat

**Affiliations:** Instituto de Patología Vegetal – Centro de Investigaciones Agropecuarias – Instituto Nacional de Tecnología Agropecuaria (IPAVE-CIAP-INTA), Camino 60 Cuadras Km 5,5 (X5020ICA), Córdoba, Argentina; Consejo Nacional de Investigaciones Científicas y Técnicas. Unidad de Fitopatología y Modelización Agrícola, Camino 60 Cuadras Km 5,5 (X5020ICA), Córdoba, Argentina; Queensland Alliance for Agriculture and Food Innovation, The University of Queensland, St. Lucia, Queensland 4072, Australia

**Author notes:** Corresponding authors: Nicolás Bejerman,; Ralf Dietzgen,; Debat Humberto.

**Keywords:** tri-segmented viruses, cytorhabdoviruses, virus taxonomy, metatranscriptomics, virus discovery, genetic diversity

## Abstract

Cytorhabdoviruses (genus *Cytorhabdovirus*, family *Rhabdoviridae*) are plant-infecting viruses with enveloped, bacilliform virions. Established members of the genus *Cytorhabdovirus* have unsegmented single-stranded negative-sense RNA genomes (ca. 10-16 kb) which encode four to ten proteins. Here, by exploring large publicly available metatranscriptomics datasets, we report the identification and genomic characterization of 93 novel viruses with genetic and evolutionary cues of cytorhabdoviruses. Strikingly, five unprecedented cytorhabdoviruses with tri-segmented genomes were also identified. This finding represents the first tri-segmented viruses in the family *Rhabdoviridae*. Interestingly, the nucleocapsid and polymerase were the only typical rhabdoviral proteins encoded by those tri-segmented viruses, whereas in three of them, a protein similar to the emaravirus (family *Fimoviridae*) silencing suppressor was found, while the other predicted proteins had no matches in any sequence databases. Genetic distance and evolutionary insights suggest that all these novel viruses may represent members of novel species. Phylogenetic analyses, of both novel and previously classified plant rhabdoviruses, provide compelling support for the division of the genus *Cytorhabdovirus* into three distinct genera. This proposed reclassification not only enhances our understanding of the evolutionary dynamics within this group of plant rhabdoviruses but also illuminates the remarkable genomic diversity they encompass. This study not only represents a significant expansion of genomics of cytorhabdoviruses that will enable future research on the evolutionary peculiarity of this genus, but also shows the plasticity in the rhabdovirus genome organization with the discovery of tri-segmented members with a unique evolutionary trajectory.

## 1. Introduction

In the current metagenomics era, the rapid discovery of novel viruses has unveiled a rich and diverse evolutionary landscape of replicating entities, that present intricate challenges in their systematic classification (Koonin et al., 2021). To address this phenomenon, diverse strategies have emerged, culminating in a comprehensive proposal for establishing a megataxonomy of the virus world (Koonin et al., 2020). However, despite extensive efforts to characterize the viral component of the biosphere, it is evident that only a minuscule fraction, likely encompassing less than one percent of the entire virosphere, has been comprehensively characterized to date (Geoghegan and Holmes, 2017; Dominguez-Huerta et al., 2023). Consequently, our understanding of the vast global virome remains limited, with its remarkable diversity and its interactions with various host organisms (Dolja et al., 2020; Edgar et al., 2022; Koonin et al., 2022; Mifsud et al., 2022). To fill this knowledge gap, researchers have used the mining of publicly available transcriptome datasets obtained through High-Throughput Sequencing (HTS) as an efficient and inexpensive strategy (Edgar et al., 2022; Bejerman et al., 2021; Bejerman et al., 2022; Debat et al., 2023). This data-driven approach to virus discovery has become increasingly valuable, given the wealth of freely available datasets within the Sequence Read Archive (SRA) maintained by the National Center for Biotechnology Information (NCBI), which is continually expanding at an extraordinary rate. This data represents a substantial, albeit still somewhat limited and potentially biased, portion of the organisms inhabiting our world, thus making the NCBI-SRA database a cost-effective and efficient resource for the identification of novel viruses (Lauber and Seitz, 2022). *Serratus* (Edgar et al., 2022) has become an invaluable and exciting tool which facilitates comprehensive data mining, thus accelerating virus sequence discovery at a pace never witnessed before. In terms of virus taxonomy, a consensus statement has emphasized the importance of incorporating viruses known solely based on metagenomic data into the official classification scheme of the International Committee on Taxonomy of Viruses (ICTV) (Simmonds et al., 2017). This recognition underscores the significance of metagenomic approaches in expanding our understanding of the global virome and adapting taxonomic frameworks to accommodate the ever-expanding diversity of viruses (Simmonds et al., 2023).

The family *Rhabdoviridae* is composed of members with negative-sense single-stranded RNA genomes which infect a broad range of hosts including plants, amphibians, fish, mammals, reptiles, insects, and other arthropods (Dietzgen et al., 2017; Walker et al., 2022). Almost all rhabdovirus genomes are unsegmented, but interestingly, plant-associated rhabdoviruses with bi-segmented genomes and a shared evolutionary history of rhabdoviruses have been included in the family in both genera *Dichorhavirus* and *Varicosavirus* (Dietzgen et al., 2017; Walker et al., 2022). *Cytorhabdovirus* is one of the genera that include plant-infecting viruses (family *Rhabdoviridae*, subfamily *Betarhabdovirinae*) (Walker et al., 2022). Most cytorhabdoviruses exhibit a genome organization characterized by the presence of six conserved canonical genes encoded in the order 3′-nucleocapsid protein (N) -phosphoprotein (P) – movement protein (P3) -matrix protein (M) -glycoprotein (G) –large polymerase (L)-5′, and up to four additional accessory genes with unknown functions, leading to diverse genome organizations (Dietzgen et al., 2020). With some exceptions, the presence and synteny of the canonical genes is strictly conserved, nevertheless, some cytorhabdoviruses lack the G gene (Bejerman et al., 2021). The viral genes are separated by conserved gene junction sequences, and the whole coding region is flanked by 3′ leader and 5′ trailer sequences that possess partially complementary ends, which could form a panhandle structure during viral replication (Dietzgen et al., 2017).

In this study, through mining of publicly available sequence data, we identified 93 novel cytorhabdoviruses including five viruses with an unprecedented tri-segmented genome, which represent the first tri-segmented genomes among rhabdoviruses. Our findings will significantly advance the taxonomical classification of cytorhabdoviruses, allowing to split this genus into three genera and shed new light on the evolutionary landscape of this group of plant rhabdoviruses.

## 2. Material and Methods

### 2.1 Identification of cytorhabdovirus-like sequences from public plant RNA-seq datasets

We analyzed the Serratus database using the Serratus Explorer tool (Edgar et al., 2022) and as query the predicted RNA-dependent RNA polymerase protein (RdRP) of cytorhabdoviruses available at the NCBI- refseq database. The SRA libraries that matched the query sequences (alignment identity > 45%; score > 10) were further explored in detail.

### 2.2 Sequence assembly and virus identification

Virus discovery was implemented as described elsewhere (Bejerman et al., 2022; Debat et al., 2023). In brief, the raw nucleotide sequence reads from each SRA experiment that matched the query sequences in the Serratus platform were downloaded from their associated NCBI BioProjects (**Tables 1-4**). The datasets were pre-processed by trimming and filtering with the Trimmomatic v0.40 tool as implemented in http://www.usadellab.org/cms/?page=trimmomatic with standard parameters. The resulting reads were assembled *de novo* with rnaSPAdes using standard parameters on the Galaxy server (https://usegalaxy.org/). The transcripts obtained from *de novo* transcriptome assembly were subjected to bulk local BLASTX searches (E-value < 1e^-5^) against cytorhabdovirus refseq protein sequences available at https://www.ncbi.nlm.nih.gov/protein?term=txid11305[Organism]. The resulting viral sequence hits of each dataset were explored in detail. Tentative virus-like contigs were curated (extended and/or confirmed) by iterative mapping of each SRA library’s filtered reads. This strategy was used to extract a subset of reads related to the query contig, used the retrieved reads from each mapping to extend the contig and then repeat the process iteratively using as query the extended sequence (Bejerman et al., 2022). The extended and polished transcripts were reassembled using Geneious v8.1.9 (Biomatters Ltd.) alignment tool with high sensitivity parameters.

### 2.3 Bioinformatics tools and analyses

#### 2.3.1 Sequence analyses

ORFs were predicted with ORFfinder (minimal ORF length 120 nt, genetic code 1, https://www.ncbi.nlm.nih.gov/orffinder/), functional domains and architecture of translated gene products were determined using InterPro (https://www.ebi.ac.uk/interpro/search/sequence-search) and the NCBI Conserved domain database - CDD v3.20 (https://www.ncbi.nlm.nih.gov/Structure/cdd/wrpsb.cgi) with e- value = 0. 1. Further, HHPred and HHBlits as implemented in https://toolkit.tuebingen.mpg.de/#/tools/ were used to complement annotation of divergent predicted proteins by hidden Markov models. Transmembrane domains were predicted using the TMHMM version 2.0 tool (http://www.cbs.dtu.dk/services/TMHMM/) and signal peptides were predicted using the SignalP version 6.0 tool (https://services.healthtech.dtu.dk/services/SignalP-6.0/). The presence of gene junction sequences flanking ORFs was also included as a criterion to determine the potential coding sequences. The predicted proteins were then subjected to NCBI-BLASTP searches against the non-redundant protein sequences (nr) database.

#### 2.3.2 Pairwise sequence identity

Percentage amino acid (aa) sequence identities of the predicted L protein of all viruses identified in this study, as well as those available in the NCBI database were calculated using SDTv1.2 (Muhire et al., 2014) based on MAFFT 7.505 (https://mafft.cbrc.jp/alignment/software) alignments with standard parameters. Virus names and abbreviations of cytorhabdoviruses already reported are shown in **Supplementary Table S1**.

#### 2.3.3 Phylogenetic analysis

Phylogenetic analysis based on the predicted L and N proteins of all plant cytorhabdoviruses, listed in **Table S1**, was done using MAFFT 7.505 with multiple aa sequence alignments using FFT-NS-i as the best-fit model. The aligned aa sequences were used as the input in Mega11 software (Tamura et al., 2021) to generate phylogenetic trees by the maximum-likelihood method (best-fit model = WAG + G + F). Local support values were computed using bootstraps with 1000 replicates. L and N proteins of selected varicosaviruses and alphanucleorhabdoviruses were used as outgroups.

## 3. Results

### 3.1. Summary of discovered viral sequences

In this study, through identification, assembly, and curation of raw NCBI-SRA reads of publicly available transcriptomic data we obtained the coding complete viral genomic sequences of 93 novel viruses with genetic and evolutionary links to cytorhabdoviruses. The phylogenetic relationships of the now significantly expanded number of known cytorhabdoviruses provide support for splitting the genus *Cytorhabdovirus* to establish three genera (**Fig. 1**) that represent distinct evolutionary lineages, which we propose to name as *Alphacytorhabdovirus* (**Table 1**), *Betacytorhabdovirus* (**Table 2**) and *Gammacytorhabdovirus* (**Table 3**). Strikingly, five unprecedented viruses with a tri-segmented genome were also identified and their full-length viral genomic sequences were assembled (**Table 4**), including the corrected full-length coding genome segments of the previously reported Picris cytorhabdovirus 1 (PiCRV1) (Rivarez et al., 2023), which had one RNA segment missing, as well as its RNA2 partially annotated.

**Figure 1.**
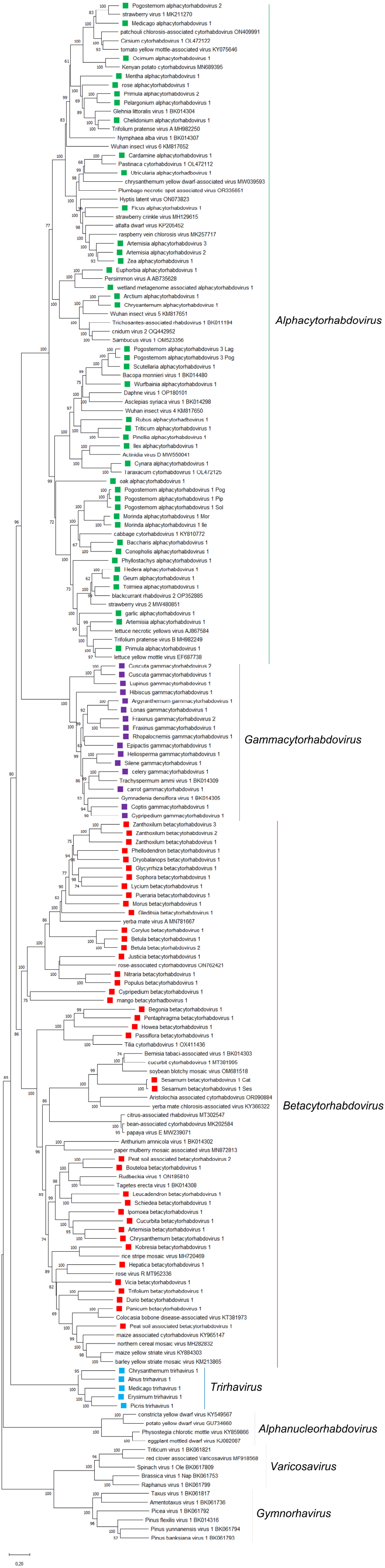
Maximum-likelihood phylogenetic tree based on amino acid sequence alignments of the complete L gene of all tri-segmented rhabdoviruses and cytorhabdoviruses reported so far and in this study constructed with the WAG + G + F model. The scale bar indicates the number of substitutions per site. Bootstrap values following 1000 replicates are given at the nodes, but only the values above 50% are shown. The viruses identified in this study are noted with green, red, violet, and blue rectangles according to proposed genus membership. Alphanucleorhabdoviruses, gymnorhaviruses and varicosaviruses were used as outgroups.

### 3.2 Genus Alphacytorhabdovirus

The full-length coding regions of 38 novel putative alphacytorhabdoviruses were assembled in this study, including three variants of the same virus associated with different plant hosts, and two host variants of two other viruses (**Table 1**). The newly identified viruses were associated with 39 plant host species and a wetland metagenome study (**Table 1**). Most of the apparent host plants are herbaceous dicots (27/39), while 6 hosts are monocots and another 6 are woody dicots (**Table 1**).

The genomic organization of the 38 novel alphacytorhabdoviruses was quite similar, with few exceptions, with six distinct genomic organizations observed (**Table 1**, **Fig.2B**). Two virus genomes have no additional accessory genes and have the genome organization 3′-N-P–P3-M-G-L-5′ (**Table 1**, **Fig. 2B**), while 14 viruses had an overlapping ORF within the P-encoding ORF, named P’, one virus had an accessory ORF between the G and L genes displaying a 3′-N-P–P3-M-G-P6-L-5′ genomic organization and 20 viruses had both those accessory ORFs (**Table 1**, **Fig. 2B**). Another virus also had two accessory ORFs, one located between the P3 and the M genes, and the other between the G and L genes, displaying a 3′-N-P–P3-P4-M-G-P7-L-5’ genomic organization (**Table 1**, **Fig. 2B**). Another newly identified virus also had two accessory ORFs, one between the G and L genes, and the other following the L gene, showing a 3′-N-P–P3-M-G-P6-L-P8-5’ genomic organization (**Table 1**, **Fig. 2B**). P4 and P8 proteins yielded no hits when BlastP searches were carried out, and no conserved domains were identified in these proteins. On the other hand, transmembrane domains were identified in each P’ protein, as well as in each protein encoded by the accessory ORF located between the G and L genes.

**Figure 2.**
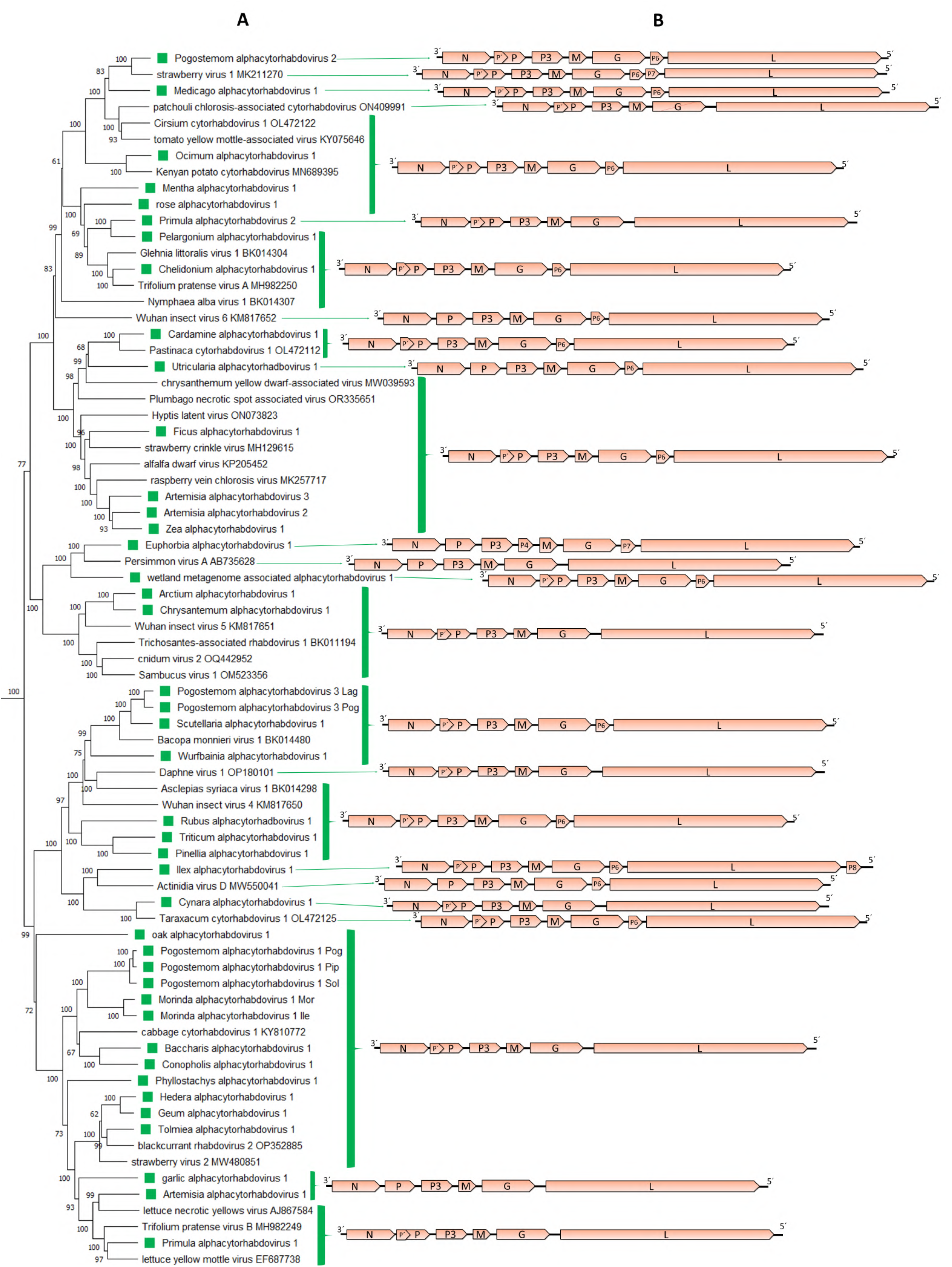
A: An inset of the maximum-likelihood phylogenetic tree shown in Fig. 1 was cropped to show those viruses included in the proposed genus *Alphacytorhabdovirus*. The viruses identified in this study are noted with green rectangles. B: genomic organization of the viral sequences used in the phylogeny.

The consensus gene junction sequences of the novel alphacytorhabdoviruses identified in our study were highly similar to those of previously reported phylogenetically related cytorhabdoviruses (**Table 5**).

Pairwise aa sequence identity values between each of the L proteins of the 38 novel viruses and those from known alphacytorhabdoviruses varied significantly, ranging from 36.16% to 85.65% (**Table S2**), while sequence identity for variants of the same virus ranged from 89.01% to 96.28% (**Table S2**). On the other hand, the highest L protein aa sequence identity with those cytorhabdoviruses proposed to be classified as betacytorhaboviruses and gammacytorhabdoviruses was 33.84% (**Table S2**).

A phylogenetic analysis based on the L protein aa sequence showed that the 38 novel viruses grouped with 33 known cytorhabdoviruses in a distinct major cluster (**Fig. 1**). Within this cluster of 71 viruses several clades could be distinguished (**Fig. 2A**). One major clade and other minor ones were composed of viruses that do not have accessory ORFs between the G and L genes (**Fig. 2**), while another clade grouped together viruses with an accessory ORF between the G and L genes (**Fig. 2**). Other clusters grouped together viruses with distinct genomic organizations (**Fig. 2**). A similar topology was observed in the phylogenetic tree based on the N protein aa sequences (**Fig. S1**).

### 3.3. Genus Betacytorhabdovirus

The full-length coding regions of 39 novel putative betacytorhabdoviruses were assembled in this study (**Table 2**), including two distinct variants of the same virus. Based on the database information, the identified viruses were associated with 36 plant host species and two peat soil metagenomes (Table 2). Interestingly, 18/36 hosts are woody dicots, while 13/36 hosts are herbaceous dicots, and the other 5 hosts are monocots (**Table 2**).

The genomic organization of the 39 novel betacytorhabdoviruses was quite diverse, with 12 distinct genomic organizations observed (**Table 2**, **Fig. 3B**). Several (12/39) viruses lack additional accessory genes and have the conserved basic genome organization 3′-N-P–P3-M-G-L-5′, but 11 of those genomes have a significantly shorter G gene (**Table 2**, **Fig. 3B**). Other viruses (16/39) had an accessory ORF between the G and L genes displaying a 3′-N-P–P3-M-G-P6-L-5′ genomic organization. One virus had an accessory ORlocated after the L gene, thus displaying a 3′-N-P–P3-M-G-L-P7-5’ genomic organization, and one virus had an accessory ORF between the P3 and M genes showing a 3′-N-P–P3-P4- M-G-L-5’ genome organization (**Table 2**, **Fig. 3B**). One virus had two accessory ORFs, one between the G and L genes, and the other after the L gene showing a 3′-N-P–P3-M-G-P6-L-P8-5’ genome organization (**Table 2**, **Fig 3.B**). Yet another virus had two accessory ORFs in the same position, however this virus lacked a discernable P3 gene, thus displaying a 3′-N-P–M-G-P5-L-P7-5’ genome organization (**Table 2**, **Fig. 3B**). Four viruses had three accessory ORFs each in their genome, located either between the G and L genes displaying a 3′-N-P–P3-M-G-P6-P7-P8-L-5’ genome organization, or two accessory ORFs between the G and L genes and another between the N and P genes showing a 3′-N-X-P–P3-M-G- P7-P8-L-5’ genome organization, or two accessory ORFs located between the P3 and the M genes, and another one after the L gene, displaying a 3′-N-P–P3-P4-M-G-L-P7-5’ genomic organization (**Table 2**, **Fig. 3B**). On the other hand, one virus appeared to only have four genes in the order 3’-N-P-P3-L-5’, while the genome of two other viruses had five genes in the order 3′-N-P–P3-M-L-5′ but lacking the G gene (**Table 2**, **Fig. 3B**). The P4 protein encoded by the virus named as Passiflora betacytorhabdovirus 1 showed no hits when BlastP searches were carried out, and no known conserved domains were identified, whereas the P4 protein encoded by Sesamum virus 1 had no hits against the database, but a transmembrane domain and a Signal peptide were predicted. No hits against the database, nor conserved domains were found in those proteins encoded by the accessory ORF located after the L gene, or by the ORFs located after the accessory ORF which encodes the P6 protein in those viruses that have more than one accessory ORF between the G and L genes. Transmembrane domains were identified in the P6 protein which is encoded by an accessory ORF located between the G and L genes in those viruses, named as P5 in Kobresia betacytorhabdovirus 1 and P7 in Justicia betacytorhabdovirus 1 and Passiflora betacytorhabdovirus 1.

**Figure 3.**
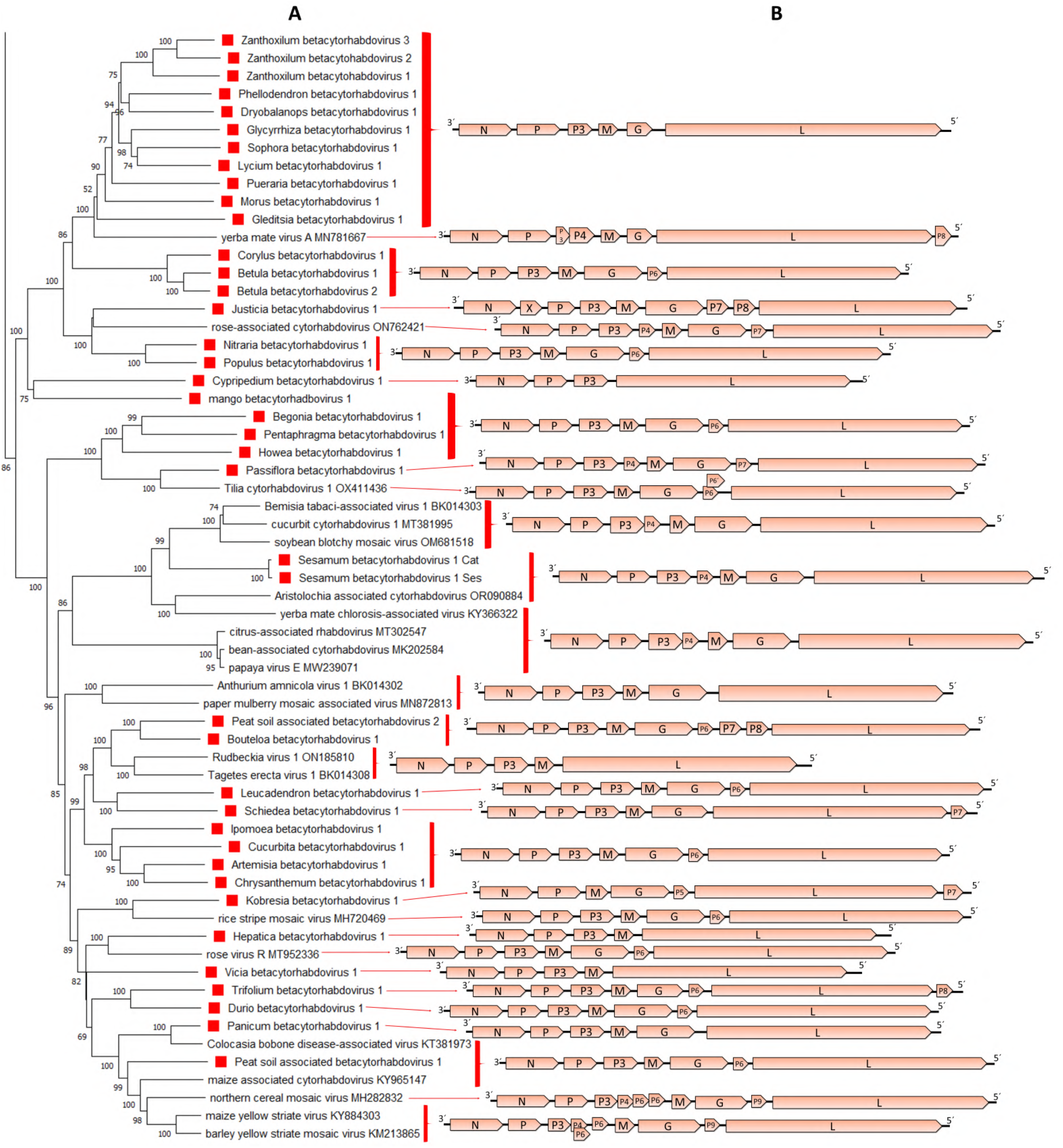
A: An inset of the maximum-likelihood phylogenetic tree shown in Fig. 1 was cropped to show those viruses included in the proposed genus *Betacytorhabdovirus*. The viruses identified in this study are noted with red rectangles. B: genomic organization of the viral sequences used in the phylogeny.

The consensus gene junction sequences among the novel betacytorhabdoviruses identified in this study and those already known, showed some variability, mainly in the length of the intergenic spacer (**Table 5**).

Pairwise aa sequence identity values between each of the L proteins of the 39 novel viruses and those from known betacytorhabdoviruses varied significantly, ranging between 27% and 80.06% (**Table S2**), while the L protein identity for variants of the same virus ranged between 93.47% and 99.29% (**Table S2**). The highest L protein aa sequence identity with those cytorhabdoviruses proposed to be classified as betacytorhaboviruses and gammacytorhabdoviruses was 33.84% (**Table S2**).

Phylogenetic analysis based on L protein aa sequences showed that the 39 novel viruses grouped with 20 known cytorhabdoviruses in a distinctive major group that we named betacytorhabdoviruses (**Fig. 1**). Within this distinct group of 59 viruses several evolutionary clades could be distinguished (**Fig. 3A**). One clade grouped together all viruses that have a short G gene and share a similar genomic organization with no additional accessory ORFs in their genomes except for Yerba mate virus A, which clustered basal to this clade (Fig. 3). Another clade grouped together viruses with two accessory genes located between the P and M genes, exemplified by Sesamum betacytorhabdovirus 1 (**Fig. 3**). Several other clades that grouped together viruses with a similar genomic organization were also observed (**Fig. 3**). However, other clusters grouped together viruses with diverse genomic organizations (**Fig. 3**). A similar topology was observed in the phylogenetic tree based on the N protein aa sequences (**Fig. S1**).

### 3.4 Genus Gammacytorhabdovirus

The full-length coding regions of 16 novel putative gammacytorhabdoviruses were assembled in this study (**Table 3**), bringing the number of potential members of this proposed genus to 18, by inclusion of two previously reported cytorhabdoviruses (**Fig. 4A**). The newly identified viruses were tentatively associated with 15 plant host species and the fungus *Hymenoscyphus fraxineus* (**Table 3**). Most of the host plants (12/15) were herbaceous dicots, while two were orchids, and one was a dicot tree (**Table 3**).

**Figure 4.**
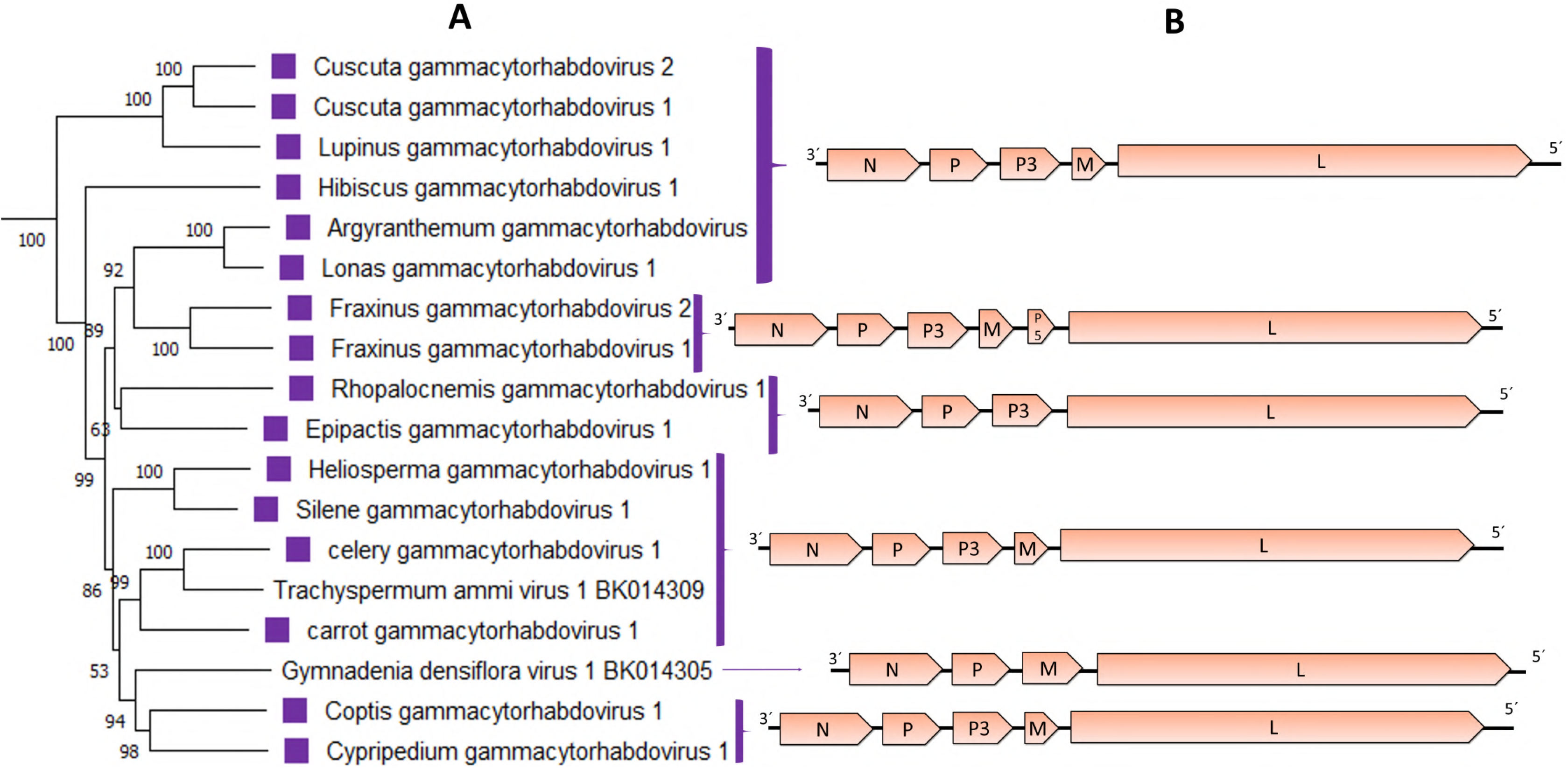
A: An inset of the maximum-likelihood phylogenetic tree shown in Fig. 1 was cropped to show those viruses included in the proposed genus *Gammacytorhabdovirus*. The viruses identified in this study are noted with violet rectangles. B: genomic organization of the viral sequences used in the phylogeny.

The genomic organization of the 16 gammacytorhabdoviruses is quite similar with only a few exceptions. A common feature of all newly identified gammacytorhabdoviruses is the absence of the G gene. Twelve viruses had five genes in the order 3′-N-P–P3-M-L-5′; while two viruses had four genes in the order 3′-N- P–P3-L-5’ and lacked the M gene. Interestingly, the two viruses associated with Fraxinus display a genomic organization 3′-N-P–P3-M-P5-L-5’, with six genes including a small ORF located between the M and L genes (**Table 3**, **Fig. 4B**). The predicted P5 protein showed no hits when BlastP searches were carried out, but one transmembrane domain was identified.

The consensus gene junction sequences of the novel gammacytorhabdoviruses identified in this study were highly similar and resembled those of the two previously reported phylogenetically related cytorhabdoviruses (**Table 5**).

Pairwise aa sequence identity values between each L protein of the 18 proposed gammacytorhabdoviruses varied significantly, ranging between 49% and 84% (**Table S2**). However, the highest sequence identity with those cytorhabdoviruses proposed to be classified as alphacytorhaboviruses and betacytorhabdoviruses was 33.4% (**Table S2**).

The phylogenetic analysis based on the L protein aa sequence showed that the 16 novel viruses grouped with two known cytorhabdoviruses in a distinct group (**Fig. 1**). Within this distinct group of 18 viruses, most of the clusters grouped together viruses with the same genome organization and/or type of hosts, such as both Fraxinus-associated viruses, the cluster that grouped the carrot, celery, and Trachyspermum- associated viruses, the cluster composed of the Heliosperma and Silene-associated viruses, or the cluster composed of the Argyranthemum and Lonas-associated viruses (**Fig. 4**). Interestingly, the orchid-associated viruses (Cypripedium, Epipactis and Gymnadenia) did not share a similar genomic organization and did not cluster together (**Fig. 4**). A similar topology was observed in the phylogenetic tree based on the N protein aa sequences (**Fig. S1**).

### 3.5 Tri-segmented rhabdoviruses

Unexpectedly, the full-length coding regions of four novel viruses that consisted of three genome segments were also assembled (**Table 4**). The best hits of the L, N, P2, P3 and P4 proteins encoded by those four tri-segmented viruses were the cognate proteins encoded by Picris cytorhabdovirus 1 (PiCRV1) (Rivarez et al., 2023). Two genome segments of this virus had previously been assembled, annotated, and deposited in GenBank (Accession # OL472127 and OL472128), but the assembled PiCRV1 N protein gene was significantly shorter than the N gene assembled for the four novel tri-segmented viruses. Consequently, we re-analyzed the SRA deposited by Rivarez et al. (2023) and we were able to extend the sequence of the N gene, but also to assemble a previously unrecognized third segment. Thus, five tri-segmented rhabdo-like viruses, subsequently referred to as trirhaviruses, were identified from the SRA data analysis. We propose to rename the Picris-associated virus as Picris trirhavirus 1.

RNA1 of all the tri-segmented viruses had one gene which encodes the L protein (**Table 4**, **Fig. 5A**). RNA2 of four of these viruses had 4 genes in the order 3’-N-P2-P3-P4-5’ while one virus has 5 genes in its RNA2 in the order 3’-N-P2-P3-P4-P5-5’ (**Table 4**, **Fig. 5A**). RNA3 of all tri-segmented viruses had 4 genes, where the first three encoded proteins, named as P6, P7 and P8, are homologous and syntenic to each other. The protein encoded at the 5′ end of segment 3 in the Chrysantheum and Medicago tri-segmented viruses is homologous to the P5 protein identified in the Alnus tri-segmented virus genome, while the proteins encoded in this position in the Erysimum, Picris and Alnus tri-segmented viruses are not homologous neither to P5, nor to each other, thus named as P9, P10 and P11, respectively (**Table 4**, **Fig. 5A**).

**Figure 5.**
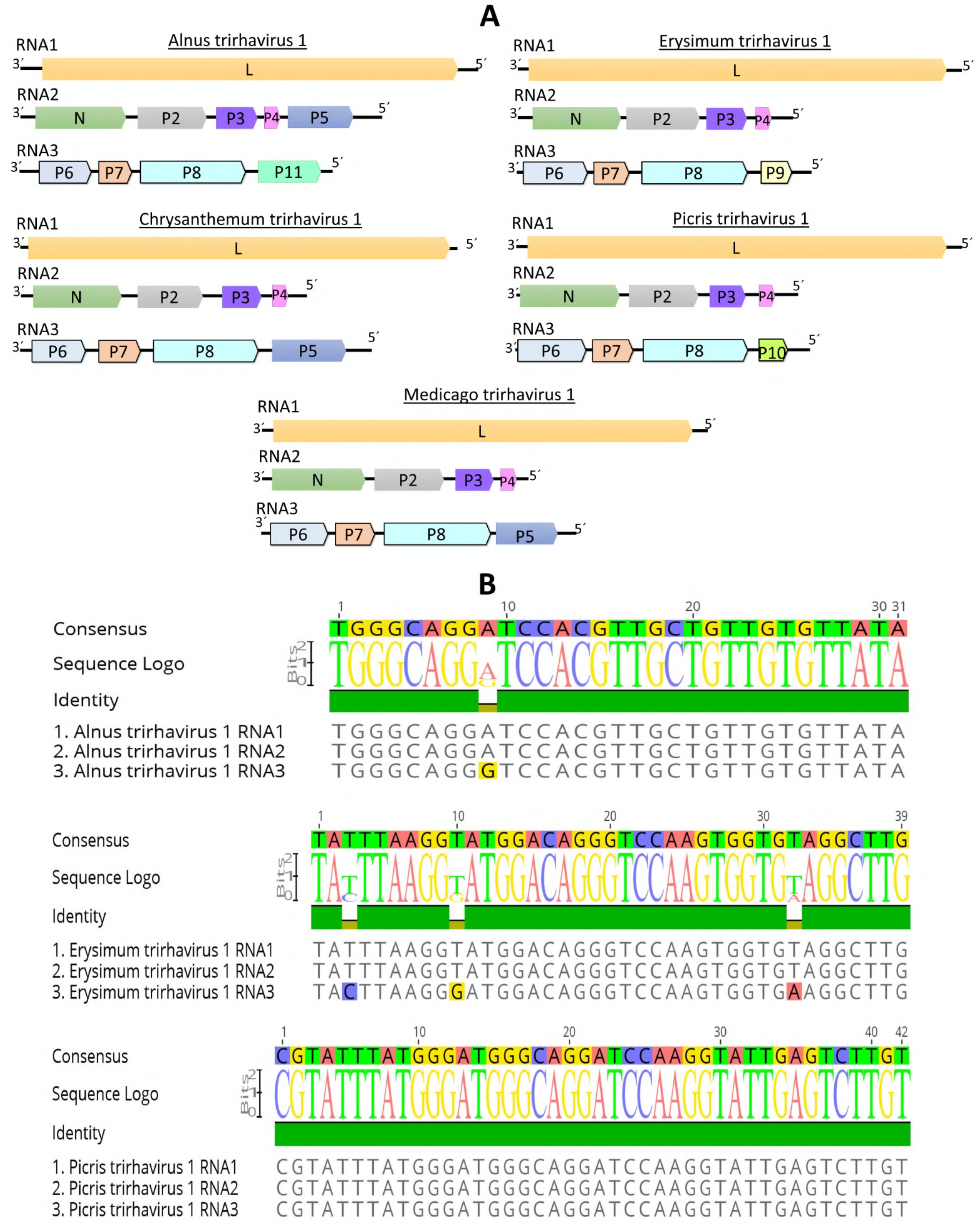
A: genomic organization of all tri-segmented rhabdoviruses identified in this study B: Alignment of the 5’trailer sequence ends of the three RNA segments of Alnus-, Erysimum- and Picris-associated viruses. The predicted coding sequences are shown in arrowed rectangles. Colors indicate protein homologies.

One interesting feature discovered when the RNA segment ends of the Alnus, Erysmum and Picris tri-segmented viruses were analyzed, is that the 30 to 40 nucleotides located at the end of the 5’trailer of each one of the segments are 99% to 100% identical (**Fig. 5B**). BlastP searches of each encoded protein showed that the L protein sequence of all tri-segmented rhabdoviruses, was more similar to the L protein of cytorhabdoviruses than to the L protein of any other rhabdovirus. On the other hand, for every tri-segmented virus, the N protein best hits were the N proteins encoded by varicosaviruses or nucleorhabdoviruses. No similarity hits were found for P2, P3, P4, P6, P7, P8, P9, P10 or P11 in databases even with relaxed parameters. Strikingly, the P5 proteins showed hits against the putative silencing suppressor protein encoded by emaraviruses (family *Fimoviridae*) (**Table 4**). A signal P was predicted in each P2 and P5 proteins, while transmembrane domains were predicted in each P4 and P8 proteins. However, no conserved domains were predicted in any of the other viral proteins.

The consensus gene junction sequences of the tri-segmented rhabdoviruses are highly similar and like those previously reported for cytorhabdoviruses proposed to be classified as alphacytorhabdoviruses (**Table 5**).

Pairwise aa sequence identity values between each the L proteins of the 5 tri-segmented viruses did not vary significantly, ranging between 55% and 66% (**Table S2**). On the other hand, the highest sequence identity with those cytorhabdoviruses proposed to be classified as alphacytorhaboviruses, betacytorhabdoviruses or gammacytorhabdoviruses was only 32% (**Table S2**). The highest sequence identity of trirhaviruses with the alpha-, beta- and gammanucleorhabdoviruses was 28.5%, and with varicosaviruses and gymnorhaviruses, the highest sequence identity was 28.6% and 27.5%, respectively. The phylogenetic analysis based on deduced L protein aa sequences placed all tri-segmented rhabdoviruses into a distinct clade which is basal to all cytorhabdoviruses (**Fig. 1**). The Alnus and Chrysantheum tri-segmented virus grouped together (**Fig. 1**), and these viruses are also the most similar in pairwise sequence identity values of their L proteins, but their RNA2 genomic organization is different (**Fig. 5A**). The second cluster included the Erysmum, Medicago and Picris tri-segmented viruses (Fig. 1), which have a similar genomic organization (**Fig.5a**). On the other hand, the phylogenetic tree based on deduced N protein aa sequences placed all tri-segmented rhabdoviruses into a distinct clade which is basal to all plant rhabdoviruses (**Fig. S1**).

## 4. Discussion

### 4.1. Discovery of novel cytorhabdo-like viruses expands their diversity and evolutionary history

In the last few years, several novel cytorhabdoviruses which do not induce visible disease symptoms have been reported in HTS studies (Bejerman et al., 2020; Belete et al., 2023; Bolus et al., 2021; Kauffmann et al., 2022; Kopke et al., 2023; Lee et al., 2022; Liu et al., 2021; Medberry et al., 2022; Petrzik et al., 2022; Safarova et al., 2022; Wu et al., 2023). Moreover, many novel cytorhabdoviruses were identified when metatranscriptomic data publicly available at the Transcriptome Shotgun Assembly (TSA) sequence databases was mined (Bejerman et al., 2021). On the other hand, the NCBI-SRA database, where many cytorhabdo-like virus sequences are likely hidden, remains significantly underexplored. This is because, traditionally, viruses were not expected to be present in sequence libraries of non-symptomatic plants. Nevertheless, the development of the Serratus tool (Edgar et al., 2022) has greatly facilitated the exploration of the SRA database, which otherwise would be tedious and time-consuming, allowing us to carry out the most extensive search to-date for cytorhabdovirus-like sequences. This substantial *in silico* directed search resulted in the identification and assembly of the full coding regions of 93 novel putative cytorhabdovirus members, representing a 1.7-fold increase of the known cytorhabdoviruses. The phylogenetic relationships, as well as the genomic features of the now expanded number of known cytorhabdoviruses, provide strong support for splitting the genus *Cytorhabdovirus* to establish three genera that we propose to name as *Alphacytorhabdovirus*, *Betacytorhabdovirus* and *Gammacytorhabdovirus*. However, the major highlight of our data mining efforts was the first-ever identification of rhabdoviruses with a tri-segmented genome. Thus, our findings clearly highlight the significance of data-driven virus discovery to increase our understanding on the genomic diversity, evolutionary trajectory, and singularity of the rhabdoviruses.

### 4.2. Proposed new genus *Alphacytorhabdovirus*

The full-length coding regions of 38 novel alphacytorhabdoviruses were assembled in this study. Most of the associated host plants were herbaceous dicots (69% of the assigned hosts), in line with previous findings as 90% of the previously identified alphacytorhabdoviruses were also associated with herbaceous dicots. Thus, these viruses likely have a host adaptation trajectory leading to preferentially infect herbaceous dicots during their evolution. The assigned hosts of six of the newly identified alphacytorhabdoviruses were monocots, and represent the first alphacytorhabdoviruses associated with monocots hosts. No apparent concordant evolutionary history with their plant hosts was observed for the monocot-infecting viruses, like what was previously reported for invertebrate and vertebrates rhabdoviruses (Geoghegan et al., 2017). Furthermore, one newly identified virus was associated with a wetland metagenome study, but even after extensive assessment of the corresponding libraries we were not able to clearly assign a host to this virus.

All but one alphacytorhabdovirus identified so far had at least the six basic plant rhabdovirus genes N, P, P3, M, G and L reported for cytorhabdoviruses (Dietzgen et al., 2020). The exception was one virus associated with the host Pogostemon, known as “patchouly chlorosis-associated cytorhabdovirus,” which was found to have a truncated G gene. It was speculated that this truncation may be linked to the fact that patchouli plants are primarily propagated vegetatively and may not require a functional G protein (Kaufmann et al., 2022). One distinctive feature of alphacytorhabdoviruses is the presence of an overlapping ORF within the P-encoding ORF, named P’ in most of their proposed members (65/71). At least one transmembrane domain was identified in each P’ protein predicted in the genomes of the alphacytorhabdoviruses assembled in this study. This is consistent with what has been previously reported for cytorhabdoviruses, where at least one transmembrane domain was identified in every P’ protein (Bejerman et al., 2021). Hence, it could be speculated that this protein serves a membrane-associated function. Nevertheless, additional research should be directed toward the functional characterization of this intriguing protein. Moreover, 42/71 alphacytorhabdoviruses have an accessory ORF between the G and L genes. The encoded small protein contains transmembrane domains, and it was speculated that it may have membrane-associated functions similar to viroporins of vertebrate rhabdoviruses (Bejerman et al., 2021). Other accessory ORFs were also detected in only two alphacytorhabdoviruses identified in this study. One of them, named P4, was located between the P3 and M genes in one virus and another one, dubbed P8, was found between the L gene and the 5’trailer. No significant hits were found for P4 or P8 when BlastP searches were carried out, and no conserved domains were identified in these proteins. Another small accessory ORF, named P7, was previously reported to be located between the P6 and L genes of strawberry virus 1 (Franova et al., 2019). Neither prediction of functional domains nor BLASTP searches against nonredundant GenBank database returned any significant hits (Franova et al., 2019). Thus, further studies should be focused on the functional characterization of the P4, P7 and P8 proteins to gain knowledge about their potential roles.

The consensus gene junction sequences among the alphacytorhabdoviruses are highly similar, likely indicating a common evolutionary history for this group of viruses. The nt sequence identity between the genomes of alphacytorhabdoviruses varied significantly ranging from 36% to 86%. This suggests that there may be still an unknown amount of “virus dark matter” within some clusters of the alphacytorhabdoviruses space worth exploring, that may contain some yet to be discovered alphacytorhabdoviruses. Moreover, the highest sequence identity with those viruses not classified as potential alphacytorhabdoviruses is very low, which is common among plant rhabdoviruses, that are characterized by a high level of diversity in both genome sequence and organization (Dietzgen et al., 2017). On the other hand, when we analyzed the diversity between variants of viruses which are likely members of the same species, the sequence identity ranged from 89% to 96%.

Among all plant rhabdoviruses studied so far, there is a strong correlation between phylogenetic relationships and vector types (Dietzgen et al., 2020). Many members grouped within the alphacytorhabdoviruses have been shown to be aphid-transmitted (Dietzgen et al., 2020; Petrzik et al., 2022), except for patchouly chlorosis-associated cytorhabdovirus, which was speculated to be vertically transmitted (Kaufmann et al., 2022). We therefore predict that the novel alphacytorhabdoviruses described here are likely aphid transmitted. In support of this, Triticum-associated virus was also found in a sequencing library of the aphid *Sitobium avenae*, thus providing some evidence for aphids as potential vectors of the newly identified alphacytorhabdoviruses.

The observed phylogenetic relationships suggest a common evolutionary history for alphacytorhabdoviruses, with four major clades observed. All viruses in the clade that includes the well-studied lettuce necrotic yellows virus do not have an accessory ORF between the G and L genes; thus, these viruses may represent the ancestral clade within the alphacytorhabdoviruses. In another clade, all members but two, had an accessory ORF between the G and L genes; therefore, these viruses may have evolved from an ancestor that already had that ORF, which is absent in the patchouly chlorosis-associated cytorhabdovirus and primula alphacytohabdovirus 2 genomes, while strawberry virus 1 acquired another accessory ORF during its evolution. In another clade, there are two major clusters, one that includes viruses that likely evolved from the ancestral ancestor, while the other cluster showed a more complex evolutionary history including members with distinct numbers of genes within their genomes. The fourth clade also showed a more complex evolutionary history because it included members with distinct genomic organizations, where many viruses acquired accessory ORFs during their evolution, mostly in the position between the G and L genes.

We propose to classify this group of evolutionary related viruses into a novel genus within the family *Rhabdoviridae*, subfamily *Betarhabdovirinae* for which we suggest the name “*Alphacytorhabdovirus*”. Based on the phylogenetic insights and the observed genetic distance of the newly identified viruses we tentatively propose 86% aa sequence identity of the L gene as threshold for species demarcation in this newly proposed genus which will include 72 members, for which the complete coding-sequence is available.

### 4.3. Proposed new genus *Betacytorhabdovirus*

The full-length coding regions of 39 novel betacytorhabdoviruses were assembled in this study. Interestingly, half of the associated hosts were woody dicots, while 35% (7/20) of the previously identified cytorhabdoviruses of this group are also associated with woody dicots. Thus, many betacytorhabdoviruses likely infect woody dicots, which may be a distinctive feature of this group of viruses. Most of the monocot-infecting betacytorhabdoviruses grouped together suggesting a shared co-divergence for these viruses. Two newly identified betacytorhabdoviruses which clustered with monocot-infecting viruses were associated with a peat soil metagenome study. Therefore, it is tempting to speculate that monocots could be associated with these viruses.

The genomic organization of the betacytorhabdoviruses is quite diverse, with 16 distinct genomic organizations discernable among its 59 putative members. Almost a quarter (14/59) of betacytorhabdoviruses lacked additional accessory genes and had at least the six basic genes N, P, P3, M, G and L reported for cytorhabdoviruses (Dietzgen et al., 2020). Nevertheless, 12 betacytorhabdoviruses had a shorter G gene. Four viruses lacked the G gene altogether, while one also lacked the M gene. The G protein was found to be essential for virus acquisition by arthropod vectors (Dietzgen et al., 2017), but it is not essential for replication and systemic movement (Wang et al., 2015). Some isolates of the betacytorhabdovirus citrus-associated rhabdovirus were recently shown to have a defective G gene, thus it was speculated that the lack of a functional G gene could provide an evolutionary advantage in fruit trees that are propagated artificially by asexual modes, such as cutting and grafting (Zhang et al., 2021). Moreover, the recently identified Rudbeckia virus 1 lacked the G gene (Lee et al., 2022), This virus was identified in Rudbeckia seeds. Thus, it was predicted to be vertically transmitted by seeds without the help of a vector which may have favored the loss of the G gene during its evolution (Lee et al., 2022). Hence, one might be inclined to speculate that viruses lacking the G gene or having a shorter G gene could potentially undergo vertical transmission. Furthermore, infections with viruses that lack the M gene have been reported to be asymptomatic (Ma et al., 2020), which is additional evidence supporting vertical transmission of the virus. Moreover, it has been shown, using a nucleorhabdovirus as model, that cooperative M-G interactions are needed for some of the functions that involve the M protein (Sun et al., 2018). Thus, perhaps in those viruses that lack the G gene, the M gene could become dispensable, and may be prone to be lost during evolution like in the Cypripedium-associated virus, which lacks both genes. Further studies should experimentally assess these conjectures. Moreover, many betacytorhabdoviruses (27/59) have an accessory ORF between the G and L genes. This small protein has transmembrane domains, and it was speculated that it may have membrane-associated functions similar to viroporins in vertebrate rhabdoviruses (Bejerman et al., 2021). Several other accessory ORFs were also identified in betacytorhabdoviruses reported in this study suggesting a complex evolutionary history where many members acquired additional ORFs during adaptation to their hosts. Four betacytorhabdoviruses had an accessory ORF between the L gene and the 5’trailer. Ten betacytorhabdoviruses had two ORFs between the P and M genes, while three others had four ORFs between these genes (Dietzgen et al., 2020). One of these ORFs encodes the putative cell-to-cell movement protein, which is named as P3 in all but the Yerba mate virus A, where this protein is named P4 (Bejerman et al., 2020). The other accessory proteins are named P4, P5 and P6. The P4 protein encoded by the newly identified Sesamum virus 1, and the one encoded by the known Bemisia tabaci associated virus, Cucurbit cytorhabdovirus 1, Yerba mate chlorosis associated virus, soybean blotchy mosaic virus, papaya virus E and Aristolochia-associated cytorhabdovirus; as well as the P5 protein encoded by barley yellow striate mosaic virus (BYSMV) and maize yellow striate virus, are small proteins (70-80aa) with a predicted transmembrane domain, suggesting a membrane association function. Indeed, the BYSMV P5 was shown to be targeted to the endoplasmic reticulum and it was suggested that the features of this protein are reminiscent of the small hydrophobic proteins of tupaia rhabdovirus (Yan et al., 2015). Two viruses had two additional accessory ORFs between the viroporin-like protein gene and the L gene, while another virus had one accessory ORF in that position. An overlapping ORF within the one encoding the viroporin-like protein was found in a cytorhabdovirus associated with the Linden tree *Tillia cordata* (Kopke et al., 2023). In the Justicia-associated virus, an accessory ORF was found between the N and P genes. An accessory ORF located in this position has been described for some alphanucleorhabdoviruses (Dietzgen et al., 2020), but Justicia-associated virus appears to be the first cytorhabdovirus with an ORF in this position. For the above accessory ORFs, except for the viroporin-like proteins and the P4/P5 proteins, BlastP results were orphans, no known signals, or domains present, and no clues towards their putative (conserved?) function were found. Thus, further studies should be focused on the functional characterization of these proteins to gain essential knowledge regarding the proteome of the accessory ORFs of betacytorhabdoviruses.

The consensus gene junction sequences of the novel and previously reported betacytorhabdoviruses showed some variability, but there appears to be a correlation with the phylogenetic relationships thus supporting a common evolutionary history for these viruses.

When we analyzed the diversity between variants of viruses which likely belong to the same species, nt sequence identity ranged from 93.5% to 99%. On the other hand, the pairwise sequence identity among betacytorhabdoviruses showed a great variation ranging between 27% and 80% which suggests that there may be many more betacytorhabdoviruses yet to be discovered. Moreover, the sequence identity with those viruses not classified as potential betacytorhabdoviruses is very low, which is a common feature among plant rhabdoviruses, that are characterized by a high level of diversity in both genome sequence and organization (Dietzgen et al., 2017). Furthermore, this high sequence diversity coupled with the distinct genomic architecture displayed by betacytorhabdoviruses, and the complex evolutionary history as shown in the phylogenetic analyses may set the foundation to further split this proposed genus in the future once additional members can be identified.

Among all plant rhabdoviruses studied so far, there is a strong correlation between phylogenetic relationships and vector types (Dietzgen et al., 2020). Some betacytorhabdoviruses have been shown to be transmitted by planthoppers, others by leafhoppers and others by whiteflies (Dietzgen et al., 2020 We therefore predict that the potential vectors of the novel betacytorhabdoviruses may be whiteflies, planthoppers, leafhoppers and likely non-aphid arthropods, like psyllids. Those betacytorhabdoviruses that lack the G gene or with a shorter G gene are likely vertically transmitted.

The phylogenetic analysis of betacytorhabdoviruses revealed several major clades suggesting a complex evolutionary history. One clade grouped together all viruses with a short G gene and without accessory genes, except for Yerba mate virus A. These viruses, with one exception, are associated with woody dicots; therefore, it is tempting to speculate that the ancestor virus was adapted to infect woody plants and that Yerba mate virus A acquired an additional ORF during its host adaptation. Another clade includes most of the viruses with two genes between the P and M genes but no accessory ORFs in other positions in their genomes likely indicating that they share the same ancestor. Certain clusters encompassed viruses that share common genomic organization, while other clusters featured viruses with unique genomic structures. An example of this diversity can be seen in the cluster that includes Cypripedium- and Mango- infecting viruses. This highlights the intricate evolutionary history of most betacytorhabdoviruses, with many of them acquiring additional genes during their evolution. Notably, these gene acquisitions were primarily concentrated between the P and M genes or between the G and L genes.

Based on the phylogenetic insights and the observed genetic distances of the newly identified viruses we tentatively propose an aa sequence identity of 82% in the L gene as threshold for species demarcation in this newly proposed genus which will include 59 members, for which the complete coding-sequence is available.

### 4.4. Proposed new genus *Gammacytorhabdovirus*

The full-length coding regions of 16 novel gammacytorhabdoviruses were assembled in this study, and most of the associated host plants were herbaceous dicots. Three viruses were linked to orchids, while two were associated with the woody tree Fraxinus but, interestingly, one of them was identified in a library of the fungal pathogen (*Hymenoscyphus fraxineus*) sampled from this woody tree.

The common feature of all 18 gammacytorhabdoviruses identified so far (16 in this study and two in Bejerman et al., 2021) is the lack of a G gene in their genome. The G gene was shown, using as a model an infectious clone of Sonchus yellow net virus, to be not essential for replication and systemic movement (Wang et al., 2015). Those two viruses associated with Fraxinus, have an additional ORF between the M and L genes, which we named P5. Interestingly, transmembrane domains were predicted for P5 suggesting a membrane-associated function for this protein that has a similar size to cytorhabdovirus viroporin-like proteins, which also have transmembrane domains (Bejerman et al., 2021). Nevertheless, no distant hits with viroporin-like proteins were found when we used HHblits on the predicted P5 protein. One previously identified gammacytorhabdovirus, associated with the orchid Gymandenia, lacks not only the G gene, but also the P3 gene. Thus, how GymDenV1 moves from cell to cell remains to be unraveled, but no cell-to-cell movement protein has either been identified in the fungi-transmitted varicosaviruses (Bejerman et al., 2021). Strikingly, two gammacytorhabdoviruses identified in this study, one associated with the orchid *Epipactis* and the other with the parasitic plant *Rhopalocnemis*, do not have M and G genes. Previous studies have assumed that the nucleocapsid core (NC) proteins N, P and L are essential for virus replication and transcription and that the M protein is required for condensation of the core during virion assembly (Dietzgen et al., 2017). M protein appears to be required for the long-distance movement of the virus within the plant (Wang et al., 2015), and an infectious clone of a plant rhabdovirus lacking the M gene displayed reduced infectivity, a vasculature-confined tissue tropism and no visible symptoms (Ma et al., 2020). Moreover, it has been shown, using a nucleorhabdovirus as model, that cooperative M-G interactions are needed for some of the functions that involve the M protein (Sun et al., 2018). Thus, it is tempting to speculate that in those viruses that lack the G gene, the M gene could be dispensable, and may have been lost during the evolution of the *Epipactis-* and *Rhopalocnemis*-associated viruses.

It has been suggested that the fungi-transmitted varicosaviruses, which do not encode a G protein (Bejerman et al., 2022), may have originated through trans-kingdom horizontal gene transfer events between fungi and plants, adapting specifically to a plant-based lifestyle (Dolja et al., 2020). The absence of the G gene, coupled with the detection of one of the recently identified gammacytorhabdoviruses in a fungal library, raises the possibility that these viruses might be transmitted by a fungal vector rather than by arthropods, as is commonly observed in viruses classified as alpha- and betacytorhabdoviruses. This serves as another distinguishing characteristic of the gammacytorhabdoviruses. Thus, further studies should focus on the potential vector and the mode of transmission of gammacytorhabdoviruses.

Another distinctive feature of gammacytorhabdoviruses is that the intergenic spacer of their gene junctions starts with an A instead of a typical G, like all other plant rhabdoviruses (Bejerman et al., 2021; 2022) suggesting a unique evolutionary history of these viruses.

The nt sequence identity among gammacytorhabdoviruses showed a high variation ranging between 49% and 84%. Moreover, the sequence identity with those viruses not classified as potential gammacytorhabdoviruses is very low, suggesting unknown gammacytorhabdovirus diversity yet to be discovered.

Interestingly, the three orchid-associated viruses (*Cypripedium*, *Gymnadenia*, and *Epipactis*) have a different genomic organization, where one virus lack the G gene, another does not encode the G and M proteins, while the third does not have the P3 and G genes. Moreover, they are not grouped together in the phylogenetic tree, thus they likely did not share a common evolutionary history. On the other hand, most of the viruses infecting herbaceous dicot hosts, as well as those associated with woody trees, clustered together according to the host family, suggesting a shared host-virus co-divergence in those clades.

We propose to classify this group of evolutionary related viruses sharing the lack of the G gene in their genomes as a distinctive feature, into a novel genus within the family *Rhabdoviridae*, subfamily *Betarhabdovirinae* for which we suggest the name “*Gammacytorhabdovirus*”. Based on the phylogenetic insights and the observed genetic distance of the newly identified viruses we tentatively propose an aa sequence identity of 85% in the L gene as threshold for species demarcation in this newly proposed genus which will include 18 members, for which the complete coding-sequences are available.

### 4.5. Tri-segmented rhabdoviruses

All rhabdoviruses identified to date have unsegmented genomes, except for the dichorhaviruses and most varicosaviruses which have bi-segmented genomes (Dietzgen et al., 2017; Bejerman et al., 2022). Unexpectedly, five novel viruses with tri-segmented genomes were identified in this study, including the corrected full-length coding genome segments of the previously reported PiCRV1 (Rivarez et al., 2023). RNA1 of all these tri-segmented viruses had only one gene that encodes the L protein, which is similar to the bi-segmented rhabdoviruses where the L protein is the only gene product of RNA1 (Dietzgen et al., 2020; Bejerman et al., 2022). RNA2 of four of the viruses has four genes, while the Alnus tri-segmented virus has five genes. Five genes are present in RNA2 of dichorhaviruses (Dietzgen et al., 2020), while three to five genes are present in RNA2 of varicosaviruses (Bejerman et al., 2022), with the N gene the only orthologous gene between them. RNA3 of all tri-segmented viruses has four genes, where the first three encoded proteins are homologous. The protein encoded at the end of this segment in the Chrysanthemum and Medicago tri-segmented viruses is homologous to P5 on RNA2 of the Alnus tri-segmented virus genome, while the proteins located in this position in the Erysimum, Picris and Alnus tri-segmented viruses are unique. This genomic organization is unique among rhabdoviruses (Dietzgen et al., 2017) and represents the first known tri-segmented rhabdovirus genomes. Other segmented negative-sense RNA viruses (NSR), belonging to the order *Bunyavirales*, have one or two genes on each RNA segment. Thus, the genomic organization of the tri-segmented rhabdoviruses identified in this study are likely distinctive among NSR viruses.

The ends of the 5’trailer region of all genome segments are conserved in the tri-segmented viruses identified in our study. A similar feature is observed in the other segmented rhabdoviruses and NSR viruses, which may be linked to RNA-dependent RNA polymerase- mediated recognition for replication. BlastP searches of the L protein encoded on RNA1 of all identified tri-segmented viruses showed that this protein is most closely related to the L Protein encoded by cytorhabdoviruses, while the best hits for the N protein were the N proteins coded by varicosaviruses or nucleorhabdoviruses. This suggests that these two proteins, which are located on different RNA segments, have distinct evolutionary histories. On the other hand, no hits were found for P2, P3, P4, P6, P7, P8, P9, P10 or P11. Strikingly, P5 showed hits against the putative RNA silencing suppressor protein encoded by emaraviruses (family *Fimoviridae*), plant viruses with segmented, linear, single-stranded, negative-sense genomes (Elbeaino et al., 2018) in the order *Bunyavirales* (Rehanek et al., 2022), while rhabdoviruses are classified in the order Mononegavirales (Walker et al., 2022). Viral RNA silencing suppressors are required for systemic infection of the plant host and the presence of these proteins suggests that the tri-segmented viruses detected here are plant-associated (Dolja et al., 2020).

A signal peptide was predicted in each P5 protein, which may be associated with its RNA silencing suppressor function, and in each P2 protein, which suggest that this protein is membrane-associated. Interestingly, a signal peptide is present in the movement protein (MP) encoded by emaraviruses (Rehanek et al., 2022), but none were identified in the MP encoded by plant rhabdoviruses (Bejerman et al., 2021), and no distant hits with any MP were found when using HHblits on the P2. Transmembrane domains were predicted in each P4 and P8 proteins suggesting a membrane-associated function for these proteins. P4 size is similar to that reported for cytorhabdovirus viroporin-like proteins, which also have transmembrane domains (Bejerman et al., 2021) while P8 size is similar to that reported for the glycoprotein encoded by plant rhabdoviruses (Bejerman et at., 2021), but no distant hit with any viroporin-like protein or glycoprotein was found using HHblits. No conserved domains were found in the other coded proteins. Thus, further studies should be focused on the functional characterization of the P2, P3, P4, P6, P7, P8, P9, P10 and P11 proteins to gain fundamental insights about the proteome of the tri-segmented viruses beyond the N, L and P5 proteins.

The novel tri-segmented viruses also resemble rhabdoviruses in possessing similar conserved gene junctions that are also highly similar to those present in the alphacytorhabdoviruses.

The pairwise aa sequence identities between the L proteins of all the tri-segmented viruses were not low at all, ranging between 55% and 66%, which may suggest that tri-segmented rhabdoviruses are evolutionarily younger than unsegmented ones.

The phylogenetic analysis based on deduced L protein aa sequences placed all tri-segmented viruses into a distinct clade within the plant rhabdoviruses that grouped with the cytorhabdoviruses rather than with varicosaviruses or nucleorhabdoviruses, whereas the phylogenetic tree based on the N protein placed the tri-segmented viruses in a clade which is basal to all plant rhabdoviruses. The complex evolutionary history of this divergent group of viruses suggests that they share a unique evolutionary history among rhabdoviruses. It is tempting to speculate that the RNA segment encoding the L protein evolved from a cytorhabdovirus ancestor, while the RNA segment encoding the N protein may have evolved from a rhabdovirus ancestor of all tri-segmented viruses, except for the Alnus-associated virus. The presence of an emaravirus-related protein in its RNA2 segment, as well as in the RNA3 segment of the Chrysanthemum- and Medicago-associated tri-segmented viruses leads us to speculate that these segments may have emerged from the recombination of a negative-sense rhabdovirus ancestor and an emaravirus. On the other hand, the RNA3 segment of the viruses from Alnus, Erysimum and Picris may have evolved from a segmented negative-sense rhabdovirus ancestor.

Taken together, these tri-segmented viruses may be taxonomically classified in a novel genus within the family *Rhabdoviridae*, subfamily *Betarhabdovirinae* for which we suggest the name “*Trirhavirus*”. Based on the phylogenetic insights and the observed genetic distance of the newly identified viruses we tentatively propose an aa sequence identity of 80% in the L gene as threshold for species demarcation in this proposed genus.

### 4.6. Strengths and Limitations of sequence discovery through data mining

As demonstrated previously by Bejerman and colleagues (2022) and in this study with the Picris- associated virus, the independent validation through re-analyzes of the NCBI-SRA raw data of viruses assembled with unexpected genomes, is important to enhance our comprehension and confidence in the genomic architecture of RNA viruses assembled via HTS data. However, the inability to revisit the original biological material for replication of results and verification of the assembled viral genome sequences is a significant weakness of the data mining approach in virus discovery. Moreover, potential issues such as contamination, low sequencing quality, spill-over, and other technical artifacts pose a risk of yielding false-positive detections, chimeric assemblies, or difficulties in accurately assigning host organisms. Therefore, researchers should be cautious when scrutinizing publicly available SRA data for virus detection. To bolster and complement such results, the acquisition of new RNAseq datasets from the predicted plant hosts is strongly recommended. Furthermore, the absence of a directed strategy for verifying genomic segment termini, such as the use of Rapid Amplification of cDNA Ends (RACE), presents challenges in determining bona fide RNA virus ends, especially considering the conserved functional and structural cues observed in rhabdoviruses (Dietzgen et al., 2017). Despite these limitations, certain aspects of our virus discovery strategy can help mitigate some of these challenges and provide additional evidence for identification. For example, when the same putative virus is consistently identified in multiple independent libraries originating from the same plant host, when there is substantial coverage of virus-related reads, when multiple RNA segments of the virus are detected within a single library, or when different viral strains are identified in plants that are closely related in terms of their evolutionary history. Nonetheless, it is essential to acknowledge that associations and detections provided in this work and other data-driven studies should be viewed as preliminary and should be complemented through subsequent studies.

## 5. Conclusions

In conclusion, this study underlines the significance of analyzing SRA public data as a valuable tool, not only for expediting the discovery of novel viruses but also for gaining insights into their evolutionary history and enhancing virus classification. Through this approach, we conducted a search for hidden cytorhabdovirus-like sequences, which significantly expanded the number of putative cytorhabdoviruses. It also allowed us to unequivocally split this group of viruses into three genera resulting in the most comprehensive cytorhabdoviruses phylogeny to date, highlighting their diversity and complex evolutionary dynamics. The major finding of our study was the first-ever identification of tri-segmented rhabdoviruses, which shows the extensive plasticity inherent to the rhabdovirus genome organization including members with unique and intriguing evolutionary trajectories. Thus, future studies should explore various unresolved aspects of these viruses, such as potential symptoms, vertical transmission, and possible vectors.

## Acknowledgments

We would like to express genuine appreciation to the producers of the original data used for this work, which are cited in **Tables 1, 2, 3 and 4**. By ensuing open science practices with accessible raw sequence data in open public repositories, they supported contributions based on secondary data analyses.

## Author Contributions

Conceptualization, N.B and H.D; data analysis, N.B and H.D; writing—original draft preparation, N.B; writing—review and editing, N.B, R.G.D and H.D. All authors have read and agreed to the published version of the manuscript.

## Institutional Review Board Statement

Not applicable for studies not involving humans or animals.

## Informed Consent Statement

Not applicable for studies not involving humans.

## Data availability statement

Nucleotide sequence data reported are available in the Third Party Annotation Section of the DDBJ/ENA/GenBank databases under the accession numbers TPA: BK064247-BK064360.

---

**Figure S1.** Maximum-likelihood phylogenetic tree based on amino acid sequence alignments of the complete N gene of all tri-segmented rhabdoviruses and cytorhabdoviruses reported so far and in this study constructed with the WAG + G + F model. The scale bar indicates the number of substitutions per site. Bootstrap values following 1000 replicates are given at the nodes, but only the values above 50% are shown. The viruses identified in this study are noted with green, red, violet, and blue rectangles according to proposed genus membership. Alphanucleorhabdoviruses, gymnorhaviruses and varicosaviruses were used as outgroups.

