## Supplementary material for "Novel tri-segmented rhabdoviruses: A data mining expedition unveils the cryptic diversity of cytorhabdoviruses": Table 1

**Table 1**. Summary of novel alphacytorhabdoviruses identified from plant RNA-seq data available on NCBI.

| **Plant host** | **Taxa/**  **family** | **Virus name/**  **Abbreviation** | **Bioproject ID/**  **Data citation** | **Length (nt)** | **Accession number** | **Protein ID** | **Length (aa)** | **Highest scoring virus- protein/*E*-value/query coverage%/identity% (Blast P)** |
| --- | --- | --- | --- | --- | --- | --- | --- | --- |
| Greater burdock  (*Arctium lappa*) | Dicot/*Asteraceae* | Arctium alphacytorhabdovirus 1/  ArcACRV1 | PRJNA598011/  Yang et al., (2022) | 12768 | BK064262 | N  P  P´  P3  M  G  L | 450  300  115  225  182  552  2097 | WhIV5-N/0.0/91/63.07  SaV1-P/1e-100/100/50.67  no hits  TrARV1-P3/6e-93/97/59.91  CnV2-M/2e-54/98/47.49  TrARV1-G/0.0/98/55.09  WhIV5-L/0.0/99/64.42 |
| Silvery wormwood  (*Artemisia argyi*) | Dicot/*Asteraceae* | Artemisia alphacytorhabdovirus 1/  ArtACRV1 | PRJNA397671/  Liu et al., (2018) | 12978 | BK064263 | N  P  P3  M  G  L | 454  315  307  184  552  2074 | LNYV-N/0.0/88/60.30  LNYV-P/9e-91/93/50.68  LNYV-P3/2e-119/96/54.52  LNYV-M/2e-48/96/45.76  LNYV-G/0.0/99/51.18  LNYV-L/0.0/99/68.25 |
| Common wormwood  (*Artemisia montana*) | Dicot/*Asteraceae* | Artemisia alphacytorhabdovirus 2/  ArtACRV2 | PRJDB8414/  Kyoto University, Japan, unpublished | 14344 | BK064264 | N  P  P´  P3  M  G  P6  L | 479  343  88  241  182  569  64  2086 | RVCV-N/1e-175/98/51.95  RCVC-P/1e-84/92/44.55  RVCV-P´/2e-10/64/49.12  RVCV-P3/2e-74/78/56.08  RVCV-M/1e-49/83/50.66  RVCV-G/0.0/92/55.51  RVCV-P6/3e-11/90/44.83  RVCV-L/0.0/99/64.72 |
| Sievers wormwood  (*Artemisia sieversiana*) | Dicot/*Asteraceae* | Artemisia alphacytorhabdovirus 3/  ArtACRV3 | PRJNA834888/  Zhang et al., (2022) | 14339 | BK064265 | N  P  P´  P3  M  G  P6  L | 476  342  103  240  185  571  64  2089 | RVCV-N/2e-171/98/50.62  RCVC-P/6e-82/93/43.69  RVCV-P´/3e-13/66/54.93  RVCV-P3/3e-75/82/52.40  RVCV-M/1e-52/82/51.97  RVCV-G/0.0/91/56.76  RVCV-P6/7e-14/100/48.44  RVCV-L/0.0/99/63.43 |
| Desert broom  (*Baccharis sarothroides*) | Dicot/*Asteraceae* | Baccharis alphacytorhabdovirus 1/  BacACRV1 | PRJNA716650/  Romero, M., UNAM, Mexico, unpublished | 13581 | BK064266 | N  P  P´  P3  M  G  L | 470  297  85  337  168  550  2074 | CCyV1-N/1e-153/87/52.68  CCyV1-P/2e-56/98/40.74  no hits  StrV2-P3/4e-120/93/56.78  CCyV1-M/5e-34/95/38.65  CCyV1-G/1e-160/92/43.14  CCyV1-L/0.0/99/59.24 |
| Large bittercress (*Cardamine amara*) | Dicot/*Brassicaceae* | Cardamine alphacytorhabdovirus 1/  CarACRV1 | PRJDB4989/  Akiyama et al., (2021) | 13209 | BK064267 | N  P  P´  P3  M  G  P6  L | 457  316  109  224  164  569  70  2092 | PaCRV1-N/0.0/99/72.35  PaCRV1-P/5e-134/63.26  PaCRV1-P´/1e-21/48.11  PaCRV1-P3/8e-122/98/75.57  PaCRV1-M/1e-89/100/73.78  PaCRV1-G/0.0/95/75.64  PaCRV1-P6/5e-26/97/63.24  PaCRV1-L/0.0/100/78.30 |
| Greater celandine  (*Chelidonium majus*) | Dicot/*Papaveraceae* | Chelidonium alphacytorhabdovirus 1/  CheACRV1 | PRJNA376854/  Zhao, L., Jinlin, China, unpublished | 12148 | BK064268 | N  P  P´  P3  M  G  P6  L | 415  325  71  200  169  552  66  2072 | TpVA-N/0.0/100/70.19  TpVA-P/2e-146/100/63.38  no hits  TpVA-P3/8e-117/98/80.71  TpVA-M/2e-73/92/69.43  TpVA-G/0.0/98/71.72  GlLV1-P6/2e-14/100/57.58  TpVA-L/0.0/100/80.41 |
| Indian chrysanthemum  (*Chrysanthemum indicum*) | Dicot/*Asteraceae* | Chrysanthemum alphacytorhabdovirus 1/  ChrACRV1 | PRJNA361213/ | 12715 | BK064269 | N  P  P´  P3  M  G  L | 448  301  140  225  19  549  2097 | WhIV5-N/0.0/99/60.22  SaV1-P/4e-100/99/51  no hits  TrARV1-P3/1e-91/97/57.92  CnV2-M/6e-50/91/45.60  SaV1-G/0.0/100/53.42  WhIV5-L/0.0/99/63.89 |
| Bear corn  (*Conopholis americana*) | Dicot/*Orobanchaceae* | Conopholis alphacytorhabdovirus 1/  ConACRV1 | PRJEB21674/  1000 Plant (1KP) Transcriptomes Initiative, Unpublished | 13083 | BK064270 | N  P  P´  P3  M  G  L | 467  299  87  328  166  546  2076 | CCyV1-N/1e-157/94/49.1  CCyV1-P/4e-70/100/42.35  no hits  StrV2-P3/3e-119/83/60.58  BCRV2-M/7e-32/93/38.06  CCyV1-G/8e-151/99/40.26  CCyV1-L/0.0/98/58.75 |
| Cardoon  (*Cynara cardunculus*) | Dicot/*Asteraceae* | Cynara alphacytorhabdovirus 1/  CynACRV1 | PRJNA590905/  Puglia et al., (2020) | 13726 | BK064271 | N  P  P´  P3  M  G  L | 472  311  132  350  179  562  2144 | TCRV1/0.0/100/80.08  TCRV1-P/3e-153/100/69.97  TCRV1-P´/1e-23/75/53.54  TCRV1-P3/0.0/99/80  TCRV1-M/3e-100/100/78.77  TCRV1-G/0.0/98/70.40  TCRV1-L/0.0/98/83.25 |
| Fischer´s spurge  (*Euphorbia fischeriana*) | Dicot/*Euphorbiaceae* | Euphorbia alphacytorhabdovirus 1/  EupACRV1 | PRJNA693977/  Zheng et al., (2022) | 13713 | BK064272 | N  P  P3  P4  M  G  P7  L | 451  330  223  130  184  559  41  2089 | PeVA-N/0.0/95/68.41  PeVA-P/2e-100/99/53.62  PeVA-P3/1e-84/100/55.61  no hits  PeVA-M/5e-57/96/48.88  PeVA-G/0.0/95/68.35  no hits  PeVA-L/0.0/99/69.3 |
| Tikoua fig  (*Ficus tikoua*) | Dicot/*Moraceae* | Ficus alphacytorhabdovirus 1/  FicACRV1 | PRJNA432314/  Bai, Y., Guiyang University, China,  unpublished | 13839 | BK064274 | N  P  P´  P3  M  G  P6  L | 458  316  84  228  182  563  69  2097 | SCV-N/1e-140/97/48.80  SCV-P/3e-52/88/36.51  no hits  SCV-P3/2e-75/100/50.43  SCV-M/7e-43/82/48.67  SCV-G/0.0/94/50.46  ADV-P6/3e-09/98/42.03  SCV-L/0.0/99/60.22 |
| Garlic  (Allium sativum) | Monocot/*Amaryllidaceae* | Garlic alphacytorhabdovirus 1/  GarACRV1 | PRJNA772184/  Liu, T., IBFC, China, unpublished | 13400 | BK064275 | N  P  P3  M  G  L | 468  298  329  174  554  2075 | LNYV-N/3e-160/93/53.17  TpVB-P/4e-53/94/34.80  StrV2-P3/5e-126/98/58.28  LNYV-M/3e-35/97/40.94  TpVB-G/5e-175/98/44.97  LYMV-L/0.0/99/58.43 |
| Herb bennet  (*Geum urbanum*) | Dicot/*Rosaceae* | Geum alphacytorhabdovirus 1/  GeuACRV1 | PRJEB23354/  Jordan et al., (2018) | 12756 | BK064276 | N  P  P´  P3  M  G  L | 459  295  98  319  177  546  2093 | StrV2-N/0.0/98/73.57  StrV2-P/7e-128/100/60.34  StrV2-P´/3e-22/98/55.67  StrV2-P3/1e-171/100/74.22  BCRV2-M/7e-78/92/68.29  BCRV2-G/0.0/96/69.57  BCRV2-L/0.0/99/72.89 |
| English ivy  (*Hedera helix*) | Dicot/*Araliaceae* | Hedera alphacytorhabdovirus 1/  HedACRV1 | PRJEB21674/  1000 Plant (1KP) Transcriptomes Initiative, Unpublished | 12588 | BK064277 | N  P  P´  P3  M  G  L | 459  295  98  319  172  546  2100 | StrV2-N/0.0/97/71.27  StrV2-P/3e-128/100/61.02  StrV2-P´/2e-22/98/53.61  StrV2-P3/2e-171/100/74.30  BCRV2-M/2e-78/99/65.50  BCRV2-G/0.0/99/68.19  BCRV2-L/0.0/98/72.95 |
| Plum-leaved holly  (*Ilex asprella*) | Dicot/*Aquifoliaceae* | Ilex alphacytorhabdovirus 1/  IleACRV1 | PRJNA736810/  Kong et al., (2022) | 14540 | BK064278 | N  P  P´  P3  M  G  P6  L  P8 | 452  333  93  370  184  553  54  2137  134 | AcCV-N/0.0/99/56.87  AcCV-P/1e-67/96/39.76  no hits  AcCV-P3/1e-130/99/52.49  AcCV-1e-33/99/37.30  AcCV-G/0.0/96/49.06  no hits  AcCV-L/0.0/98/60.77  no hits |
| Lucerne  (*Medicago sativa*) | Dicot/*Fabaceae* | Medicago alphacytorhabdovirus 1/  MedACRV1 | PRJNA644634/  Liu et al., (2022) | 13586 | BK064279 | N  P  P´  P3  M  G  P6  L | 431  360  64  224  191  547  80  2085 | StrV1-N/0.0/97/60.48  StrV1-1e-73/98/39.12  StrV1-P´/2e-13/100/54.69  StrV1-P3/7e-81/99/52.47  StrV1-M/5e-43/83/44.65  StrV1-G/0.0/99/60.44  StrV1-P6/6e-11/76/45.9  StrV1-L/0.0/99/68.21 |
| Horse mint  (*Mentha longifolia*) | Dicot/*Lamiaceae* | Mentha alphacytorhabdovirus 1/  MenACRV1 | PRJNA779119/  Wang, B., Shaoguan University, unpublished | 12387 | BK064280 | N  P  P´  P3  M  G  P6  L | 422  316  101  195  163  551  67  2072 | TpVA-N/9e-163/100/54.14  TpVA-/7e-45/96/32.18  no hits  TpVA-P3/6e-68/97/55.38  GlLV1-M/6e38/95/45.16  TpVA-G0.0/97/55.84  GlLV1-P6/4e-07/100/50.75  TpVA-L/0.0/99/62.64 |
| Indian mulberry  (*Morinda officinalis*) | Dicot/*Rubiaceae* | Morinda alphacytorhabdovirus 1_Mor/MorACRV1_Mor | PRJNA717096/  Li et al., (2023) | 13023 | BK064281 | N  P  P´  P3  M  G  L | 463  300  81  346  172  548  2075 | CCyV1-N/0.0/99/56.87  CCyV1-P/7e-68/98/42.71  no hits  BCRV2-P3/1e-115/76/59.7  CCyV1-M/7e-29/92/37.58  CCyV1-G/6e-171/93/45.14  CCyV1-L/0.0/99/59.16 |
| Chinese holly  (*Ilex cornuta*) | Dicot/*Aquifoliaceae* | Morindaalphacytorhabdovirus 1_Ile/MorACRV1_Ile | PRJNA399054/  Zeng et al.,(2019) | 12876 | BK064282 | N  P  P´  P3  M  G  L | 463  300  85  346  184  549  2075 | CCyV1-N/0.0/99/56.29  CCyV1-P/1e-66/98/43.39  no hits  BCRV2-P3/2e-116/76/60.08  CCyV1-M/2e-27/88/36.31  CCyV1-G/7e-167/98/43.57  CCyV1-L/0.0/99/59.16 |
| Oak  (*Quercus robur*) | Dicot/*Fagaceae* | Oak alphacytorhabdovirus 1/  OakACRV1 | PRJNA322128/  Broberg et al., (2018) | 12817 | BK064283 | N  P  P´  P3  M  G  L | 453  304  104  376  176  543  2078 | LYMV-N/2e-109/93/40.47  LYMV-P/1e-20/88/29.58  no hits  AscSyV1-P3/2e-46/66/35.97  CCyV1-M/3e-06/59/28.57  TrARV1-G/3e-7/98/30.16  BCRV2-L/0.0/99/45.57 |
| Holy basil  (*Ocimum tenuiflorum*) | Dicot/*Lamiaceae* | Ocimum alphacytorhabdovirus 1/  OciACRV1 | PRJNA251328/  Upadhyay et al., (2015) | 12478 | BK064284 | N  P  P´  P3  M  G  P6  L | 425  327  93  205  155  563  60  2078 | KePCyV-N/0.0/99/76.65  KePCyV-P/1e-116/99/54.41  no hits  KePCyV-P3/1e-105/99/70.53  KePCyV-M/5e-58/98/56.21  KePCyV-G/0.0/96/70.46  StrV1-P6/2e-08/83/50  KePCyV-L/0.0/99/77.36 |
| Scented pelargonium  (*Pelargonium* X *hybrid*) | Dicot/*Geraniaceae* | Pelargonium alphacytorhabdovirus 1/  PelACRV1 | PRJNA883637/  Saint-Marcoux, D., Lyon University, France, unpublished | 12332 | BK064285 | N  P  P´  P3  M  G  P6  L | 413  328  84  203  170  552  58  2073 | TpVA-N/1e-164/98/56.23  TpVA-P/2e-53/97/36.25  no hits  TpVA-P3/2e-91/96/68.02  TpVA.M/5e-37/90/44.81  TpVA-G/0.0/95/60.71  no hits  TpVA-L/0.0/99/63.7 |
| Moso bamboo  (*Phyllostachys edulis*) | Monocot/*Poaceae* | Phyllostachys alphacytorhabdovirus 1/  PhyACRV1 | PRJNA350353/  Huang et al., (2022) | 12947 | BK064286 | N  P  P´  P3  M  G  L | 455  296  83  328  193  544  2070 | LNYV-N/2e-138/97/46.55  LNYV-P/2e-44/93/37.28  no hits  StrV2-P3/7e-140/96/59.81  LNYV-M/6e-28/90/34.27  StrV2-G/8e-144/90/40.93  BCRV2-L/0.0/99/57.21 |
| Peltate green dragon  (*Pinellia peltata*) | Monocot/*Araceae* | Pinellia alphacytorhabdovirus 1/  PinACRV1 | PRJNA623739/  Wang et al., (2020) | 13438 | BK064287 | N  P  P´  P3  M  G  P6  L | 479  305  87  373  170  568  61  2106 | AscSyV1-N/4e-131/90/44.24  DV1-P/8e-47/98/36.75  no hits  AscSyV1-P3/9e-105/89/47.51  TCRV1-M/1e-05/88/31.21  WhIV4/2e-134/95/36.73  no hits  WhIV4/0.0/99/48.13 |
| Patchouli  (*Pogostemom cablin*) | Dicot/*Lamiaceae* | Pogostemom alphacytorhabdovirus 1_Pog/  PogACRV1_Pog | PRJNA660501/  Yan et al., (2021) | 13171 | BK064288 | N  P  P´  P3  M  G  L | 462  300  81  347  179  557  2070 | CCyV1-N/0.0/99/55.17  CCyV1-P/1e-69/96/40.79  no hits  StrV2-P3/3e-116/77/61.94  CCyV1-M/3e-38/92/36.14  CCyV1-G/1e-160/93/40.92  CCyV1-L/0.0/58.03 |
| Black pepper  (*Piper nigrum*) | Dicot/*Piperaceae* | Pogostemom alphacytorhabdovirus 1_ Pip/  PogACRV1_Pip | PRJNA580359/  Dantu et al., (2021) | 13063 | BK064289 | N  P  P´  P3  M  G  L | 462  300  81  347  179  557  2070 | CCyV1-N/0.0/99/54.96  CCyV1-P/3e-67/99/40.79  no hits  StrV2-P3/3e-116/77/62.31  CCyV1-M/5e-39/87/39.74  CCyV1-G/7e-164/91/42.88  CCyV1-L/0.0/58.75 |
| Tropical soda apple  (*Solanum viarum*) | Dicot/*Solanaceae* | Pogostemom alphacytorhabdovirus 1_Sol/  PogACRV1_Sol | PRJNA666394/  Pandey et al., (2018) | 13138 | BK064290 | N  P  P´  P3  M  G  L | 462  300  116  351  179  558  2070 | CCyV1-N/3e-179/99/54.84  CCyV1-P/5e-63/99/39.79  no hits  StrV2-P3/4e-115/76/61.57  CCyV1-M/4e-40/87/40.38  CCyV1-G/2e-157/95/40.98  CCyV1-L/0.0/58.31 |
| Patchouli  (*Pogostemom cablin*) | Dicot/*Lamiaceae* | Pogostemom alphacytorhabdovirus 2/  PogACRV2 | PRJNA660501/  Yan et al., (2021) | 13209 | BK064291 | N  P  P´  P3  M  G  P6  L | 421  359  64  224  179  549  69  2083 | StrV1-N/0.0/98/78.71  StrV1-P/8e-164/100/66.12  StrV1-P´/9e-25/100/73.44  StrV1-P3/4e-134/100/81.7  StrV1-M/2e-82/99/65.36  StrV1-G/0.0/100/74.05  StrV1-P6/2e-31/100/72.46  StrV1-L/0.0/99/82.4 |
| Patchouli  (*Pogostemom cablin*) | Dicot/*Lamiaceae* | Pogostemom alphacytorhabdovirus 3_Pog/  PogACRV3_Pog | PRJNA511937/  Tang et al., (2019) | 13252 | BK064292 | N  P  P´  P3  M  G  P6  L | 449  293  86  353  165  544  71  2110 | BmV1/0.0/99/71.05  BmV1-P/1e-127/100/64.85  BmV1-P´/5e-20/100/52.33  BmV1-P3/0.0/96/72.14  BmV1-M/1e-53/93/51.3  BmV1-G/0.0/95/71.43  no hits  BmV1-L/0.0/99/72.52 |
| Crepe myrtle  (*Lagerstroemia indica*) | Dicot/*Lythraceae* | Pogostemom alphacytorhabdovirus 3_Lag/  PogACRV3_Lag | PRJNA32094/  Zhang et al., (2014) | 13149 | BK064293 | N  P  P´  P3  M  G  P6  L | 449  294  86  353  180  544  71  2108 | BmV1/0.0/99/72.20  BmV1-P/1e-128/100/63.61  BmV1-P´/3e-23/100/55.81  BmV1-P3/0.0/96/72.14  BmV1-M/4e-55/83/55.63  BmV1-G/0.0/95/70.10  no hits  BmV1-L/0.0/99/72.04 |
| Candelabra primrose  (*Primula chungensis*) | Dicot/*Primulaceae* | Primula alphacytorhabdovrus1/  PriACRV1 | PRJNA616180/  Wang, X., BI, Kunming, China, unpublished | 12953 | BK064294 | N  P  P´  P3  M  G  L | 450  307  103  311  174  549  2066 | LYMV-N/0.0/99/72.1  LYMV-P/2e-127/98/61.26  no hits  LYMV-P3/2e-161/100/70.74  LYMV-M/2e-68/98/56.4  LYMV-G/0.0/98/60.19  LYMV-L/0.0/100/73.91 |
| Glory primrose  (*Primula oreodoxa*) | Dicot/*Primulaceae* | Primula alphacytorhabdovirus 2/  PriACRV2 | PRJNA544868/  Zhao et al., (2019) | 12146 | BK064295 | N  P  P´  P3  M  G  L | 414  327  82  201  167  559  2072 | TpVA-N/1e-160/98/54.57  GlLV1-P/2e-51/99/33.63  no hits  TpVA-P3/4e-86/98/64.5  GlLV1-M/5e-42/91/43.79  GlLV1-G/0.0/94/58.87  TpVA-L0.0/99/64.58 |
| Beach rose  (*Rosa rugosa*) | Dicot/*Rosaceae* | Rose  alphacytorhabdovirus 1/  RosACRV1 | PRJNA498442/  Shen et al., (2019) | 12601 | BK064296 | N  P  P´  P3  M  G  P6  L | 425  313  80  167  172  593  67  2068 | TpVA-N/7e-152/98/51.78  TpVA-P/3e-56/95/37.38  no hits  GlLV1-P3/4e-67/97/57.83  TpVA-M/5e-27/100/34.48  GlLV1-G/0.0/94/50.45  GlLV1-P6/1e-04/100/41.79  TpVA-L/0.0/99/64.85 |
| Korean bramble  (*Rubus coreanus*) | Dicot/*Rosaceae* | Rubus  alphacytorhabdovirus 1/  RubACRV1 | PRJNA401210/  Chen et al., (2018) | 14682 | BK064297 | N  P  P´  P3  M  G  P6  L | 474  297  93  366  186  573  64  2109 | DV1-N/91/3e-108/40.14  DV1-P/2e-38/98/33.22  DV1-P´/0.029/88/32.56  BmV1-P3/2e-106/93/45.45  DV1-M/5e-10/81/28.1  WhIV4-G/6e-154/93/41.2  no hits  WhIV4-L/0.0/99/47.44 |
| Barbed skullcap  (*Scutellaria barbata*) | *Dicot/Lamiaceae* | Scutellaria  alphacytorhabdovirus 1/  ScuACRV1 | PRJNA653305/  Li et al., (2023) | 13187 | BK064298 | N  P  P´  P3  M  G  P6  L | 447  295  86  350  165  547  75  2103 | BmV1-N/0.0/99/68.6  BmV1-P/4e-121/100/61.36  BmV1-P´/3e-24/100/53.49  BmV1-P3/0.0/98/72.46  BmV1-M/9e-52/93/50.65  BmV1-G/0.0/95/70.86  no hits  BmV1-0.0/99/71.44 |
| Piggyback plant  (*Tolmiea menziesii*) | Dicot/*Saxifragaceae* | Tolmiea alphacytorhabdovirus 1/  TolACRV1 | PRJNA507776/  Visger et al., (2019) | 12746 | BK064299 | N  P  P´  P3  M  G  L | 460  295  102  333  182  545  2091 | StrV2-N/0.0/97/67.11  BCRV2-P/4e-130/100/62.03  StrV2-P´/3e-19/96/43.88  BCRV2-P3/7e-166/94/72.01  BCRV2-M/3e-78/92/67.86  BCRV2-G/0.0/96/73.14  BCRV2-L/0.0/99/77.05 |
| Wheat  (*Triticum aestivum*) | Monocot/*Poaceae* | Triticum alphacytorhabdovirus 1/  TriACRV1 | PRJNA577739/  Li, Y., Hebei, China, unpublished |  | BK064300 | N  P  P´  P3  M  G  P6  L | 474  315  87  345  190  561  55  2106 | AscSyV1-N/1e-134/93/44.02  DV1-P/2e-47/97/36.81  DV1-P´/0.021/97/34.12  AscSyV1-P3/5e-109/90/48.96  BmV1-M/3e-07/98/27.15  DV1-G/6e-130/86/39.64  no hits  WhIV4-L/0.0/98/48.4 |
| Long-leaved bladderwort  (*Utricularia longifolia*) | Dicot/*Lentibulariaceae* | Utricularia alphacytorhabdovirus 1/  UtrACRV1 | PRJNA354080/  Tang, C., Nanjing University, China, unpublished | 13017 | BK064301 | N  P  P3  M  G  P6  L | 454  324  217  203  571  63  2089 | PaCRV1-N/4e-157/98/49.89  PaCRV1-P/6e-57/100/35.17  PaCRV1-P3/2e73/98/52.47  PaCRV1-M/1e-46/80/41.1  PaCRV1-G/0.0/98/48.23  no hits  PaCRV1-L/0.0/98/59.1 |
| Wetland metagenome | - | Wetland metagenome associated alphacytorhabdovirus 1/  WMaACRV1 | PRJNA338276/  Angle et al., (2017) | 12726 | BK064302 | N  P  P´  P3  M  G  P6  L | 445  301  99  219  172  551  52  2093 | PeVA-N/4e-174/89/57.39  PeVA-P/1e-71/99/43.93  no hits  PeVA-P3/1e-60/98/44.39  PeVA-M/2e-45/99/43.93  PeVA-G/0.0/95/53.86  no hits  PeVA-L/0.0/99/62.45 |
| Malabar cardamon  (*Wurfbainia villosa*) | Monocot/*Zingiberaceae* | Wurfbainia alphacytorhabdovirus 1/  WurACRV1 | PRJNA471573/  Wang, H., Guangzhou, China, unpublished | 13348 | BK064303 | N  P  P´  P3  M  G  P6  L | 465  297  93  356  187  546  85  2116 | BmV1-N/0.0/98/60.87  BmV1-P/6e-81/99/46.49  BmV1-P´/1e-09/89/43.37  BmV1-P3/8e-143/97/56.32  BmV1-M/1e-19/81/32.68  BmV1-G/0.0/96/57.47  no hits  BmV1-L/0.0/99/56.5 |
| Maize  (*Zea mays*) | Monocot/*Poaceae* | Zea alphacytorhabdovirus 1/  ZeaACRV1 | PRJNA543910/  Wang, J., Anhui, China, unpublished | 14358 | BK064304 | N  P  P´  P3  M  G  P6  L | 477  329  93  242  181  573  64  2085 | RVCV-N/0.0/97/53.45  RVCV-P/3e-90/96/46.5  RVCV-P´/1e-05/50/48.94  RVCV-P36e-78/82/54.5  RVCV-M/6e-48/96/48.28  RVCV-G/0.0/95/55.21  RVCV-P6/5e-11/100/42.19  RVCV-L/0.0/99/65.42 |

* Acronyms of best hits are listed in Supp. Table S1.
