## Supplementary material for "Novel tri-segmented rhabdoviruses: A data mining expedition unveils the cryptic diversity of cytorhabdoviruses": Table 2

**Table 2**. Summary of novel betacytorhabdoviruses identified in plant RNA-seq data available on NCBI.

| **Plant host** | **Taxa/**  **family** | **Virus name/**  **Abbreviation** | **Bioproject ID/**  **Data citation** | **Length (nt)** | **Accession number** | **Protein ID** | **Length (aa)** | **Highest scoring virus- protein/*E*-value/query coverage%/identity% (Blast P)** |
| --- | --- | --- | --- | --- | --- | --- | --- | --- |
| Rock wormwood  (*Artemisia rupestris*) | Dicot/*Asteraceae* | Artemisia betacytorhabdovirus 1/  ArtBCRV1 | PRJNA730219/  Zhou et al., (2021) | 13426 | BK064305 | N  P  P3  M  G  P6  L | 484  340  201  202  524  84  2076 | NCMV-N/2e-41/94/28.1  no hits  RudV1-P3/3e-09/62/26.72  no hits  PpVE-G/7e-24/89/22.67  no hits  RudV1-L/0.0/99/42.91 |
| Zip begonia  (*Begonia conchifolia*) | Dicot/*Begoniaceae* | Begonia betacytorhabdovirus 1/  BegBCRV1 | PRJEB26711/  Emelianova et al., (2021) | 13838 | BK064306 | N  P  P3  M  G  P6  L | 447  299  181  206  572  68  2161 | TiCRV1-N/5e-52/62/34.21  TiCRV1-P/2e-19/97/28.04  no hits  TiCRV1-M/7e-10/83/23.12  TiCRV1-G/2e-70/85/27.93  no hits  TiCRV1-L/0.0/99/41.72 |
| White birch  (*Betula pendula*) | Dicot/*Betulaceae* | Betula betacytorhabdovirus 1/  BetBCRV1 | PRJEB29260/  Alonso-Seera et al., (2019) | 14744 | BK064307 | N  P  P3  M  G  P6  L | 449  485  195  196  556  138  2242 | RaCV-N/8e-69/90/34.24  no hits  RaCV-P4/6e-19/91/23.46  RaCV-M/1e-04/83/23.78  RaCV-G/3e-59/92/27.08  no hits  RaCV-L/0.0/92/40.01 |
| Himalayan birch  (*Betula utilis*) | Dicot/*Betulaceae* | Betula betacytorhabdovirus 2/  BetBCRV2 | PRJNA638802/  Kumar, N., CSIR, India, unpublished | 15147 | BK064308 | N  P  P3  M  G  P6  L | 443  470  196  193  551  138  2246 | RaCV-N/2e-78/92/36.01  no hits  RaCV-P4/3e-22/89/28.41  RaCV-M/4e-08/86/25.68  RaCV-G/2e-53/87/27.44  no hits  RaCV-L/0.0/94/39.58 |
| Buffalo grass  (*Bouteloa dactyloides*) | Monocot/*Poaceae* | Bouteloa betacytorhabdovirus 1/  BouBCRV1 | PRJNA297834/  Amaradasa and Amudsen (2016) | 14127 | BK064309 | N  P  P3  M  G  P6  P7  P8  L | 452  381  195  199  508  72  255  190  2070 | RudV1-N/2e-65/95/32.07  RVR-P/1e-07/17/33.33  RudV1-P3/9e-36/89/39.13  RudV1-M/5e-28/84/34.32  NCMV-G/5e-21/94/23.43  no hits  no hits  no hits  RudV1-L/0.0/99/49.04 |
| Hardy garden mum (*Chrysanthemum morifolium*) | Dicot/*Asteraceae* | Chrysanthemum betacytorhabdovirus 1/  ChrBCRV1 | PRJNA397042/  Yue et al., (2018) | 13309 | BK064310 | N  P  P3  M  G  P6  L | 450  333  198  206  511  86  2075 | MaCyV-N/1e-45/92/30.07  RVR-P/0.007/26/29.55  TaEV1-P3/3e-11/67/29.93  no hits  PpVe-G/7e-22/69/24.66  no hits  RudV1-0.0/99/42.21 |
| Siberian hazelnut  (*Corylus heterophylla*) | Dicot/*Betulaceae* | Corylus betacytorhabdovirus 1/  CorBCRV1 | PRJNA899668/  Sun, J, Lianoning, China, unpublished | 15228 | BK064311 | N  P  P3  M  G  P6  L | 461  479  197  201  555  141  2257 | RaCV/7e-64/93/31.96  no hits  YmVA-P4/4e-14/71/28.97  RaCV-M/4e-04/87/21.59  PpVE-G/3e-73/94/27.58  no hits  RaCV-L/0.0/89/40.21 |
| Buffalo gourd  (*Cucurbita foetidissima*) | Dicot/*Cucurbitaceae* | Cucurbita betacytorhabdovirus 1/  CucBCRV1 | PRJNA473174/  Sun, University of California, USA, unpublished | 12969 | BK064312 | N  P  P3  M  G  P6  L | 450  302  195  209  514  79  2079 | MaCyV-N/1e-47/94/29.88  no hits  RudV1-P3/3e-08/59/31.03  RudV1-M/2e-06/71/27.81  RSMV-G/5e-22/93/23.35  no hits  RudV1-L/0-0/99/41.7 |
| Slipper orchid (*Cypripedium flavum*) | Monocot/*Orchidaceae* | Cypripedium betacytorhabdovirus 1/  CypBCRV1 | PRJNA479379/  Guo et al., (2018) | 9958 | BK064313 | N  P  P3  L | 430  280  198  2114 | MYSV-N/1e-38/56/36  no hits  no hits  RaCV-L/0.0/91/33.81 |
| Keladan  (*Dryobalanops oblongifolia*) | Dicot/*Dipterocarpaceae* | Dryobalanops betacytorhabdovirus 1/  DryBCRV1 | PRJDB8182/  Ng et al., (2021) | 14393 | BK064314 | N  P  P3  M  G  L | 495  598  234  273  261  2259 | YmVA-N/1e-95/98/35.8  YmVA-P/6e-07/23/32  YmVa-P4/1e-29/68/38.04  no hits  no hits  YmVA-L/0.0/98/42.34 |
| Durian  (*Durio zibethinus*) | Dicot/*Malvaceae* | Durio betacytorhabdovirus 1/  DurBCRV1 | PRJNA400310/  Teh et al., (2017) | 12791 | BK064315 | N  P  P3  M  G  P6  L | 434  318  193  174  539  62  2057 | NCMV-N/6e-52/97/33.03  no hits  PMuMaV-P3/8e-13/68/28.79  no hits  RSMV-G/6e-34/84/24.74  no hits  MYSV-L/0.0/98/45.16 |
| Littleleaf honey locust  (*Gleditsia microphylla*) | Dicot/*Fabaceae* | Gleditsia betacytorhabdovirus 1/  GleBCRV1 | PRJNA848854/  Yang et al., (2022) | 13339 | BK064316 | N  P  P3  M  G  L | 483  454  240  251  162  2240 | YmVA-N/9e86/99/33.61  no hits  YmVA-P4/6e-37/69/37.35  no hits  no hits  YmVA-L/0.0/99/40.15 |
| Chinese licorice  (*Glycyrrhiza inflata*) | Dicot/ *Fabaceae* | Glycyrrhiza betacytorhabdovirus 1/  GlyBCRV1 | PRJNA574093/  Yang et al., (2022) | 14755 | BK064317 | N  P  P3  M  G  L | 493  500  238  280  169  2264 | YmVA-N/8e-95/96/37.89  YmVA-P/3e-25/19/57.14  YmVA-P4/2e-39/76/37.57  no hits  no hits  YmVA-L/0.0/98/42.05 |
| Pennywort  (*Hepatica nobilis*) | Dicot/*Ranunculaceae* | Hepatica betacytorhabdovirus 1/  HepBCRV1 | PRJDB6630/  Nodai Genome Research Center, Japan, unpublished | 10440 | BK064318 | N  P  P3  M  L | 432  384  183  162  2074 | RVR-N/2e-77/92/35.78  CBDaV-P/6e-07/52/24.63  RVR-P3/9e-13/70/27.91  no hits  RVR-L/0.0/99/48.03 |
| Kentia palm  (*Howea forsteriana*) | Monocot/*Arecaceae* | Howea betacytorhabdovirus 1/  HowBCRV1 | PRJNA244607/  Dunning et al., (2016) | 13727 | BK064319 | N  P  P3  M  G  P6  L | 447  301  173  211  557  69  2145 | TiCRV1-N/7e-46/53/36.86  TiCRV1-P/6e-10/95/24.76  TiCRV1-P3/5e-11/81/25.53  TiCRV1-M/1e-10/91/24.37  TiCRV1-G/1e-58/92/26.15  no hits  TiCRV1-L/0.0/99/40.22 |
| Sweet potato  (*Ipomoea batatas*) | Dicot/*Convolvulaceae* | Ipomoea betacytorhabdovirus 1/  IpoBCRV1 | PRJNA626066/  Read, ARC, SouthAfrica, unpublished | 12811 | BK064320 | N  P  P3  M  G  P6  L | 448  327  196  210  533  101  2071 | NCMV-N/2e-55/93/32.62  RVR-P/0.001/25/26.76  RudV1-P3/1e-06/56/30  RudV1-M/0.015/70/24.68  RSMV-G/1e-22/86/22.76  no hits  TaEV1-1/0.0/99/41.46 |
| Malabar nut  (*Justicia adhatoda*) | Dicot/*Acanthaceae* | Justicia betacytorhabdovirus 1/  JusBCRV1 | PRJNA842169/  Padmanabhan et al., (2022) | 15957 | BK064321 | N  X  P  P3  M  G  P7  P8  L | 463  180  408  198  199  574  150  140  2236 | RaCV-N/3e-89/99/37.26  no hits  no hits  RaCV-P4/4e-31/95/31.94  RaCV-M/1e-08/83/23.7  RaCV-G/8e-96/89/32.44  RaCV-P7/1e-06/82/29.27  no hits  RaCV/0.0/96/45.34 |
| Royle´s sedge  (*Kobresia royleana*) | Monocot/*Cyperaceae* | Kobresia betacytorhabdovirus 1/  KobBCRV1 | PRJNA588660/  Qu, G., Lhasa, China, unpublished | 14255 | BK064322 | N  P  M  G  P5  L  P7 | 542  550  175  545  89  2098  106 | RSMV-N/7e-91/97/36.17  RSMV-P/8e-07/17/34.38  RSMV-M/0.003/88/26.28  RSMV-G/1e-124/94/39.24  no hits  RsMV-L/0.0/99/52.39  no hits |
| Plate-seed conebush  (*Leucadendron platyspermum*) | Dicot/*Proteaceae* | Leucadendron betacytorhabdovirus 1/  LeuBCRV1 | PRJEB45774/  Scharmann et al., (2021) | 12698 | BK064323 | N  P  P3  M  G  P6  L | 445  408  192  188  533  63  2081 | NCMV-N/1e-115/93/43.97  NCMV-P/2e-32/72/31.76  NCMV-P3/3e-09/76/26.35  TaEV1-M/8e-08/86/26.83  RSMV-G/9e-34/94/26.07  no hits  RudV1-L/0.0/99/42.01 |
| Black goji  (*Lycium ruthenicum*) | Dicot/*Solanaceae* | Lycium betacytorhabdovirus 1/  LycBCRV1 | PRJNA505629/  Zhao et al., (2020) | 14855 | BK064324 | N  P  P3  M  G  L | 504  515  239  286  208  2260 | YmVA-N/2e-78/92/35.16  YmVA-P/1e-20/22/45.76  YmVA-P4/2e-37/95/34.06  YmVA-M/0.003/67/23.35  no hits  YmVA-L/0.0/98/41.21 |
| Mango  (*Mangifera indica*) | Dicot/*Anacardiaceae* | Mango betacytorhabdovirus 1/  ManBCRV1 | PRJNA487154/  Wang et al., (2020) | 13826 | BK064325 | N  P  P3  M  G  P6  L | 477  359  167  195  577  95  2148 | YmVA-N/8e-35/84/26.25  no hits  no hits  no hits  RVR-G/7e-20/78/22.41  no hits  RaCV-L/0.0/94/35.18 |
| White mulberry  (*Morus alba*) | Dicot/*Moraceae* | Morus betacytorhabdovirus 1/  MorBCRV1 | PRJNA597172/  Jiao et al., (2020) | 15904 | BK064326 | N  P  P3  M  G  L | 502  617  231  263  215  2260 | YmVA-N/8e-99/91/36.15  YmVA-P/5e-32/74/29.3  YmVA-P4/5e-39/84/37.44  no hits  no hits  YmVA-L/0.0/98/43.14 |
| Nitre bush  (*Nitraria tangutorum*) | Dicot/*Nitrariaceae* | Nitraria betacytorhabdovirus 1/  NitBCRV1 | PRJNA686177/  Chen et al., (2021) | 15520 | BK064327 | N  P  P3  M  G  P6  L | 433  544  189  187  583  153  2246 | RaCV-N/2e-121/99/42.96  RaCV-P/2e-14/51/27.97  RaCV-P4/1e-38/95/37.57  RaCV-M/2e-21/95/30.56  RaCV-G/6e-98/86/34.9  RaCV-P7/3e-09/59/32.97  RaCV-L/0.0/99/44.89 |
| Hall´s panicgrass  (*Panicum hallii*) | Monocot/*Poaceae* | Panicum betacytorhabdovirus 1/  PanBCRV1 | PRJNA306692/  Lovell et al., (2016) | 12136 | BK064328 | N  P  P3  M  G  L | 418  280  195  172  505  2068 | CBDaV-N/0.0/99/72.9  CBDaV-P/2e-128/100/66.07  CBDaV-P3/3e-104/94/75.14  CBDaV-M/1e-83/98/68.05  CBDaV-G/0.0/100/66.14  CBDaV-L/0.0/100/75.87 |
| Blue passionflower  (*Passiflora caerulea*) | Dicot/*Passifloraceae* | Passiflora betacytorhabdovirus 1/  PasBCRV1 | PRJEB21674/  1000 Plant (1KP) Transcriptomes Initiative, Unpublished | 13471 | BK064329 | N  P  P3  P4  M  G  P7  L | 500  317  188  69  209  541  72  2138 | TiCRV1-N/8e-130/92/44.49  TiCRV1-P/6e-35/89/32.77  TiCRV1-P3/1e-22/80/29.14  no hits  TiCRV1-M/5e-48/85/47.19  TiCRV1-G/4e-160/90/44.85  no hits  TiCRV1-L/0.0/99/62.66 |
| Peat soil | - | Peat soil associated betacytorhabdovirus 1/  PSaBCRV1 | PRJNA412438/  Hausmann et al., (2019) | 12663 | BK064330 | N  P  P3  M  G  P6  L | 436  315  186  169  505  56  2083 | MYSV-N/5e-96/99/38.79  NCMV-P/2e-26/86/31.62  BYSMV-P3/1e-21/78/36.99  MYSV-M/7e-09/94/24.07  MaCyV-G/2e-75/95/31.43  no hits  MaCyV-L/0.0/99/53.75 |
| Peat soil | - | Peat soil associated betacytorhabdovirus 2/  PSaBCRV2 | PRJNA570134/  JGI, USA, unpublished | 14865 | BK064331 | N  P P3  M  G  P6  P7  P8  L | 457  397  194  199  516  66  276  182  2066 | RudV1-N/2e-65/88/33.41  BYSMV-P/4e-09/26/35.51  RudV1-P3/9e-29/78/36.84  RudV1-M/2e-21/88/31.64  PpVe-G/8e-35/82/25.61  no hits  no hits  no hits  RudV1-L/0.0/99/50.63 |
| *Pentaphragma spicatum* | Dicot/ *Pentaphragmataceae* | Pentaphragma betacytorhabdovirus 1/  PenBCRV1 | PRJNA636634/  Zhang et al., (2020) | 12983 | BK064332 | N  P  P3  M  G  P6  L | 446  288  178  197  550  63  2160 | TiCRV1-N/2e-50/57/34.91  TiCRV1-P/8e-17/87/27.21  no hits  TiCRV1-M/0.008/84/25.44  TiCRV1-G/3e-76/91/30.18  no hits  TiCRV1-L/0.0/99/41.12 |
| Amur cork tree  (*Phellodendron amurense*) | Dicot/*Rutaceae* | Phellodendron betacytorhabdovirus 1/  PheBCRV1 | PRJNA817294/  Li et al., (2022) | 14292 | BK064333 | N  P  P3  M  G  L | 488  541  241  294  251  2258 | YmVA-N/1e-85/97/34.43  YmVA-P/3e-40/61/34.47  YmVA-P4/7e-30/78/32.28  no hits  no hits  YmVA-L/0.0/98/41.42 |
| Desert poplar  (*Populus pruinosa*) | Dicot/*Salicaceae* | Populus betacytorhabdovirus 1/  PopBCRV1 | PRJNA354971/  Yu, L., Lanzhou, China, unpublished | 15094 | BK064334 | N  P  P3  M  G  P6  L | 432  569  188  201  586  150  2246 | RaCV-N/8e-118/99/42.73  RaCV-P/6e-10/50/22.37  RaCV-P4/1e-33/90/34.71  RaCV-M/1e-17/85/26.59  RaCV-G/2e-102/92/32.84  RaCV-P7/2e-10/78/31.15  RaCV-L/0.0/99/45.75 |
| Kudzu  (*Pueraria montana*) | Dicot/*Fabaceae* | Pueraria betacytorhabdovirus 1/  PueBCRV1 | PRJNA515956/  He et al., (2019) | 13614 | BK064335 | N  P  P3  M  G  L | 481  338  230  255  166  2254 | YmVA-N/1e-69/97/33.74  YmVA-P/8e-11/75/41.35/  YmVA-P4/3e-20/58/36.57  no hits  no hits  YmVA-L/0.0/98/39.88 |
| Sesame  (*Sesamum indicum*) | Dicot/*Pedaliaceae* | Sesamum  betacytorhabdovirus 1_Ses/  SesBCRV1_Ses | PRJNA644139/  Dutta et al., (2022) | 13565 | BK064336 | N  P  P3  P4  M  G  L | 439  340  183  76  224  575  2113 | CuCV1-N/5e-72/95/34.95  YmCaV-P/9e-15/58/30.10  SbBMV-P3/2e-28/73/41.18  no hits  CuCV1-M/1e-23/75/30.59  YmCaV-G/2e-100/84/35.74  CuCV1-L/0.0/99/48.07 |
| Madagascar periwinkle (*Catharanthus roseus*) | Dicot/*Apocynaceae* | Sesamum  betacytorhabdovirus 1_Cat/  SesBCRV1_Cat | PRJNA246273/  Verma et al., (2014) | 13497 | BK064337 | N  P  P3  P4  M  G  L | 440  340  183  76  224  575  2113 | CuCV1-N/1e-71/95/35.33  YmCaV-P/9e-15/58/30.10  SbBMV-P3/9e-28/73/41.18  no hits  CuCV1-M/7e-24/75/30.59  YmCaV-G/3e-100/84/35.95  CuCV1-L/0.0/99/48.02 |
| *Schiedea pentandra* | Dicot/*Caryophyllaceae* | Schiedea betacytorhabdovirus 1/  SchBCRV1 | PRJNA491458  Nevado et al., (2019) | 12964 | BK064338 | N  P  P3  M  G  L  P7 | 439  379  206  182  524  2062  114 | MYSV-N/3e-55/93/33.01  NCMV-P/6e-95/98/42.89  RVR-P3/8e-10/62/30.47  no hits  RSMV-G/1e-32/95/24.86  RudV1-L/0.0/99/43.63  no hits |
| Japanese pagoda tree  (*Sophora japonica*) | Dicot/*Fabaceae* | Sophora betacytorhabdovirus 1/  SopBCRV1 | PRJNA797104/  Song et al., (2023) | 13767 | BK064339 | N  P  P3  M  G  L | 501  493  241  283  137  2255 | YmVA-N/3e-94/91/36.54  YmVA-P/2e-37/72/31.27  YmVA-P4/1e-39/93/35.29  YmVA-M/0.001/61/24.57  no hits  YmVA-L/0.0/97/41.79 |
| Red clover  (*Trifolium pratense*) | Dicot/*Fabaceae* | Trifolium betacytorhabdovirus 1/  TriBCRV1 | PRJNA561285/  Herber et al., (2021) | 13511 | BK064340 | N  P  P3  M  G  P6  L  P8 | 429  372  218  176  529  72  2069  175 | BYSMV-N/4e-46/90/34.43  CBDaV-P/6e-06/30/26.55  PMuMaV-P3/5e-19/66/31.65  AntAmV1-M/4e-04/66/25.42  RSMV-G/4e-37/91/25.2  no hits  BYSMV-L/0.0/99/45.15  no hits |
| Broad bean  (*Vicia faba*) | Dicot/ *Fabaceae* | Vicia betacytorhabdovirus 1/  VicBCRV1 | PRJNA591424/  Yang et al., (2020) | 12101 | BK064341 | N  P  P3  M  L | 434  441  186  164  2099 | MaCyV-N/3e-60/95/31.13  RVR-P/0.035/16/30.99  RVR-P3/8e-22/75/34.04  no hits  CBDaV-L/0.0/98/44.27 |
| Japanese prickly ash  (*Zanthoxilum ailanthoides*) | Dicot/*Rutaceae* | Zanthoxilum betacytorhabdovirus 1/  ZanBCRV1 | PRJNA656412/  Chuang et al., (2021) | 16669 | BK064342 | N  P  P3  M  G  L | 488  627  243  276  278  2278 | YmVA-N/1e-88/92/33.98  YmVA-P/2e-25/22/47.18  YmVA-P4/9e-26/70/33.53  no hits  no hits  YmVA-L/0.0/98/41.48 |
| Japanese prickly ash  (*Zanthoxilum ailanthoides*) | Dicot/*Rutaceae* | Zanthoxilum betacytorhabdovirus 2/  ZanBCRV2 | PRJNA656412/  Chuang et al., (2021) | 15584 | BK064343 | N  P  P3  M  G  L | 492  579  242  270  281  2280 | YmVA-N/8e-88/91/35.01  YmVA-P/6e-37/61/33.33  YmVA-P4/8e-27/68/37.35  no hits  no hits  YmVA-L/0.0/98/40.79 |
| Japanese prickly ash  (*Zanthoxilum ailanthoides*) | Dicot/*Rutaceae* | Zanthoxilum betacytorhabdovirus 3/  ZanBCRV3 | PRJNA656412/  Chuang et al., (2021) | 16283 | BK064344 | N  P  P3  M  G  L | 492  578  242  272  283  2282 | YmVA-N/3e-88/98/34.2  YmVA-P/6e-27/17/57.84  YmVA-P4/2e-24/80/32.82  no hits  no hits  YmVA-L/0.0/99/41.24 |

* Acronyms of best hits are listed in Supp. Table S1.
