## Supplementary material for "Novel tri-segmented rhabdoviruses: A data mining expedition unveils the cryptic diversity of cytorhabdoviruses": Table 3

**Table 3**. Summary of novel gammacytorhabdoviruses identified from plant RNA-seq data available on NCBI.

| **Plant host** | **Taxa/**  **family** | **Virus name/**  **Abbreviation** | **Bioproject ID/**  **Data citation** | **Length (nt)** | **Accession number** | **Protein ID** | **Length (aa)** | **Highest scoring virus- protein/*E*-value/query coverage%/identity% (Blast P)** |
| --- | --- | --- | --- | --- | --- | --- | --- | --- |
| Teide marguerite (*Argyranthemum tenerifae*) | Dicot/*Asteraceae* | Argyranthemum gammacytorhabdovirus 1/  ArgGCRV1 | PRJNA491458/  Nevado et al., (2019) | 10801 | BK064345 | N  P  P3  M  L | 450  297  231  176  2068 | GymDenV1-N/1e-98/94/41.31  no hits  TrAV1-P3/7e-28/96/29.91  GymDenV1-M/1e-24/88/33.97  GymDenV1-L/0.0/99/55.15 |
| carrot (*Daucus carota*) | Dicot/*Apiaceae* | Daucus gammacytorhabdovirus 1/  DauGCRV1 | PRJNA745346/  Chakrabarti, S., CSIR-IICB, unpublished | 11730 | BK064346 | N  P  P3  M  L | 459  328  230  201  2069 | TrAV1-N/2e-133/95/47.42  TrAV1-P/2e-36/99/30.65  TrAV1-P3/4e-52/95/39.73  GymDenV1-M/2e-23/91/33.33  TrAV1-L/0.0/99/64.34 |
| celery (*Apium graveolens*) | Dicot/*Apiaceae* | Apium gammacytorhabdovirus1/  ApiGCRV1 | PRJNA543957/  Zhang et al., (2019) | 12008 | BK064347 | N  P  P3  M  L | 455  325  233  197  2069 | TrAV1-N/4e-173/94/57.83  TrAV1-P/2e-81/87/47.44  TrAV1-P3/4e-81/94/53-95  TrAV1-M/4e-61/94/51.87  TrAV1-L/0.0/100/72.5 |
| Chinese goldthread  (*Coptis chinensis*) | Dicot/*Ranunculaceae* | Coptis gammacytorhabdovirus 1/  CopGCRV1 | PRJNA361017/  Chen et al., (2017) | 11214 | BK064348 | N  P  P3  M  L | 437  286  227  187  2069 | GynDenV1-N/5e-118/99/42.6  TrAV1-P/2e-30/97/28.52  TrAV1-P3/1e-47/93/35.81  GymDenV1-M/2e-40/93/41.95  TrAV1-L/0.0/99/61.74 |
| Bigseed alfalfa dodder (*Cuscuta indecora*) | Dicot/*Convolvulaceae* | Cuscuta gammacytorhabdovirus 1/  CusGCRV1 | PRJNA543296/  Johnson et al., (2019) | 10772 | BK064349 | N  P  P3  M  L | 429  301  220  196  2054 | TrAV1-N/7e-108/96/43.68  GymDenV1-P/3e-21/92/29.87  TrAV1-P3/3e-16/97/25.23  GymDenV1-M/6e-13/80/32.1  GymDenV1-L/0.0/99/50.17 |
| Nevada dodder  (*Cuscuta nevadensis*) | Dicot/*Convolvulaceae* | Cuscuta gammacytorhabdovirus 2/  CusGCRV2 | PRJNA561399/  Frangione, E., Canada, unpublished | 10700 | BK064350 | N  P  P3  M  L | 429  302  220  188  2054 | TrAV1-N/1e-106/96/42.49  GymDenV1-P/9e-20/83/27.97  TrAV1-P3/2e-17/95/27.78  GymDenV1-M/6e-17/85/34.15  GymDenV1-L/0.0/99/50.85 |
| Slipper orchid (*Cypripedium flavum*) | Monocot/*Orchidaceae* | Cypripedium gammacytorhabdovirus 1/  CypGCRV1 | PRJNA479379/  Guo et al., (2018) | 10872 | BK064351 | N  P  P3  M  L | 437  283  228  213  2069 | GymDenV1-N/2e-117/96/45.5  GymDenV1-P/2e-46/96/35.1  TrAV1-P3/4e-36/94/34.86  GymDenV1-M/4e-38/83/37.64  GymDenV1-L/0.0/99/60.34 |
| Violet helleborine  (*Epipactis purpurata*) | Monocot/*Orchidaceae* | Epipactis gammacytorhabdovirus 1/  EpiGCRV1 | PRJNA450088/  Lallemand et al., (2019) | 11001 | BK064352 | N  P  P3  L | 452  300  225  2064 | GymDenV1-N/2e-102/86/42.36  GymDenV1-P/4e-21/93/26.51  TrAV1-P3/2e-26/64/34.72  GymDenV1-L/0.0/99/57.83 |
| Common ash  (*Fraxinus excelsior*) | Dicot/*Oleaceae* | Fraxinus gammacytorhabdovirus 1/  FraGCRV1 | PRJEB4958/  Sollars et al., (2017) | 11521 | BK064353 | N  P  P3  M  P5  L | 443  284  224  184  65  2068 | TrAV1-1e-96/94/41.96  GymDenV1-P/1e-26/94/28.81  TrAV1-P3/7e-29/97/29.41  GymDenV1-M/2e-30/85/38.22  no hits  GymDenV1-L0.0/99/55.78 |
| Ash dieback (*Hymenoscyphus fraxineus*) | - | Fraxinus gammacytorhabdovirus 2/  FraGCRV2 | PRJEB7998/  Rallapalli et al., (2015) | 11737 | BK064354 | N  P  P3  M  P5  L | 439  285  224  187  55  2068 | GymDenV1-2e-101/89/40.61  GymDenV1-P/8e-39/94/30.51  TrAV1-P3/6e-33/96/34.84  GymDenV1-M/7e-31/86/36.65  no hits  GymDenV1-L0.0/99/56 |
| Dwarf heliosperma (*Heliosperma pusillum*) | Dicot/*Caryophyllaceae* | Heliosperma gammacytorhabdovirus 1/  HelGCRV1 | PRJNA760819/  Szukala et al., (2023) | 11579 | BK064355 | N  P  P3  M  L | 436  308  221  206  2063 | GymDenV1-N/3e-102/90/41.65  GymDenV1-P/2e-30/84/31.9  TrAV1-P3/1e-27/95/31.63  GymDenV1-M/5e-29/83/35.67  GymDenV1-L/0.0/99/58.31 |
| Kenaf  (*Hibiscus cannabinus*) | Dicot/*Malvaceae* | Hibiscus gammacytorhabdovirus 1/  HibGCRV1 | PRJNA602109/  Chen et al., (2014) | 11079 | BK064356 | N  P  P3  M  L | 458  391  221  194  2063 | GymDenV1-N/3e-77/88/35.39  GymDenV1-P/6e-08/62/25.99  TrAV1-P3/2e-16/78/26.92  GymDenV1-M/9e-12/79/26.45  TrAV1-L/0.0/99/53.86 |
| Golden ageratum  (*Lonas annua*) | Dicot/*Asteraceae* | Lonas gammacytorhabdovirus 1/  LonGCRV1 | PRJNA371565/  Jayasena et al., (2017) | 11920 | BK064357 | N  P  P3  M  L | 450  297  231  176  2068 | GymDenV1-N/5e-106/88/44.75  GymDenV1-P/1e-20/94/26.56  TrAV1-P3/3e-24/95/27.6  GymDenV1-M/5e-24/88/30.77  GymDenV1-L/0.0/99/55.40 |
| Mantano river lupine  (*Lupinus mantaroensis*) | Dicot/*Fabaceae* | Lupinus gammacytorhabdovirus 1/  LupGCRV1 | PRJNA318864/  Nevado et al., (2016) | 11196 | BK064358 | N  P  P3  M  L | 430  314  221  189  2057 | TrAV1-N/8e-105/98/40.95  GymDenV1-P/9e-20/85/25.91  TrAV1-P3/1e-16/76/30.41  GymDenV1-M/1e-08/84/27.16  TrAV1-L/0.0/99/51.47 |
| Stinkhorn clubhead  (*Rhopalocnemis phalloides*) | Dicot/*Balanophoraceae* | Rhopalocnemis  gammacytorhabdovirus 1/  RhoGCRV1 | PRJNA737177/  Yu et al., (2022) | 11024 | BK064359 | N  P  P3  L | 469  305  231  2071 | GymDenV1-N/6e-110/86/43.06  GymDenV1-P/1e-17/84/26.16  TrAV1-P3/7e-25/92/29.17  GymDenV1-L/0.0/99/55.86 |
| Bladder campion  (*Silene vulgaris*) | Dicot/*Caryophyllaceae* | Silene gammacytorhabdovirus 1/  SilGCRV1 | PRJNA104951/  Baloun et al., (2014) | 11500 | BK064360 | N  P  P3  M  L | 435  311  221  209  2066 | GymDenV1-N/6e-107/89/43.83  GymDenV1-P/3e-35/88/29.14  TrAV1-P3/9e-33/97/31.96  GymDenV1-M/5e-31/80/36.09  GymDenV1-L/0.0/99/58.04 |

* Acronyms of best hits are listed in Supp. Table S1.
