## Supplementary material for "Novel tri-segmented rhabdoviruses: A data mining expedition unveils the cryptic diversity of cytorhabdoviruses": Table 4

**Table 4**. Summary of trirhaviruses identified from plant RNA-seq data available on NCBI, including the reannotation of Picris cytorhabdovirus 1 sequence.

| **Plant host** | **Taxa/**  **family** | **Virus name/**  **Abbreviation** | **Bioproject ID/**  **Data citation** | **RNA segment/**  **Length (nt)** | **Accession number** | **Protein ID** | **Length (aa)** | **Highest scoring virus- protein/*E*-value/query coverage%/identity% (Blast P)** |
| --- | --- | --- | --- | --- | --- | --- | --- | --- |
| Red alder  (*Alnus rubra*) | *Dicot/Betulaceae* | Alnus  trirhavirus 1/  AlTRV1 | PRJNA691057/  Bell, C., NCGR, USA,unpublished | RNA1 6699  RNA2 5289    RNA3 4586 | BK064247  BK064248  BK064249 | L  N  P2  P3  P4  P5  P6  P7  P8  P11 | 2043  442  341  201  72  312  260  165  515  289 | PiCRV1-L/0.0/99/55  PiCRV1-N/6e-62/78/34.72  PiCRV1-40kDa/2e-148/99/62.28  PiCRV1-21kDA/5e-21/89/29.61  PiCRV1-8kDa/1e-15/100/52.78  PCLSaV-P5/4e-26/53/34.94  no hits  no hits  no hits  no hits |
| Hardy garden mum (*Chrysanthemum morifolium*) | Dicot/*Asteraceae* | Chrysanthemum trirhavirus 1/  ChTRV1 | PRJNA510496/  Shen R, China, unpublished | RNA1 6332  RNA2 4222    RNA3 5133 | BK064250  BK064251  BK064252 | L  N  P2  P3  P4  P6  P7  P8  P5 | 2047  441  348  189  72  265  194  528  354 | PiCRV1-L/0.0/99/58.11  PiCRV1-N/8e-73/77/37.29  PiCRV1-40kDa/7e-154/98/63.19  PiCRV1-21kDA/1e-31/90/35.84  PiCRV1-8kDa/3e-13/100/47.22  no hits  no hits  no hits  PCLSaV-P5/1e-25/47/37.43 |
| Sierra Nevada wallflower  (*Erysimum nevadense*) | Dicot/*Brassicaceae* | Erysimum trirhavirus1/  EryTRV1 | PRJNA473238/  Osuna-Mascaró et al., (2023) | RNA1 6524  RNA2 3989    RNA3 4307 | BK064253  BK064254  BK064255 | L  N  P2  P3  P4  P6  P7  P8  P9 | 2039  441  346  198  94  316  199  509  143 | PiCRV1-L/0.0/99/66.22  PiCRV1-N/4e-111/79/45.98  PiCRV1-40kDa/2e-163/99/63.48  PiCRV1-21kDA/1e-36/85/39.18  PiCRV1-8kDa/3e-22/76/62.5  no hits  no hits  no hits  no hits |
| Lucerne  (*Medicago sativa*) | Dicot/*Fabaceae* | Medicago  trirhavirus 1/  MeTRV1 | PRJNA667169/Medina et al., (2021)  and  PRJNA535257/  JGI, USA, unpublished | RNA1 6495  RNA2 3851    RNA3 4565 | BK064256  BK064257  BK064258 | L  N  P2  P3  P4  P6  P7  P8  P5 | 2040  445  343  183  72  274  189  514  303 | PiCRV1-L/0.0/99/60.28  PiCRV1-N/2e-113/77/48.47  PiCRV1-40kDa/3e-149/97/60.90  PiCRV1-21kDA/2e-26/96/32.78  PiCRV1-8kDa/1e-18/100/62.5  no hits  no hits  no hits  PCLSaV-P5/1e-14/52/33.33 |
| Bristly ox-tongue  (*Picris echioides*) | Dicot/*Asteraceae* | Picris trirhavirus 1/  PiTRV1 | PRJNA772045/  Rivarez et al., (2023) | RNA1 6530  RNA2 4091    RNA3 4259 | BK064259  BK064269  BK064261 | L  N  P2  P3  P4  P6  P7  P8  P10 | 2043  495  345  184  72  331  199  505  148 | PiCRV1-L/0.0/100/100  PiCRV1-N/0.0/72/100  PiCRV1-40kDa/0.0/100/100  PiCRV1-21kDA/5e-134/100/100  PiCRV1-8kDa/2e-42/100/100  no hits  no hits  no hits  no hits |

* Acronyms of best hits are listed in Supp. Table S1.
