## Supplementary material for "Novel tri-segmented rhabdoviruses: A data mining expedition unveils the cryptic diversity of cytorhabdoviruses": Table 5

**Table 5**. Consensus conserved plant rhabdovirus gene junction sequences

| **Proposed Genus** | **Virus*** | **3´end mRNA** | **intergenic spacer** | **5´end mRNA** |
| --- | --- | --- | --- | --- |
| *Alphacytorhabdovirus* | ArcACRV1 | AAUUAUUUU | GAU | CUU |
|  | ArtACRV1 | AAUUCUUUU | GA(U)_n_ | CNN |
|  | ArtACRV2 | AAUUAUUUU | GA(U)_n_ | CNN |
|  | ArtACRV3 | AAUUAUUUU | GA(U)_n_ | CNU |
|  | BacACRV1 | AAUUCUUUU | GA(U)_n_ | CNC |
|  | CarACRV1 | AAUUAUUUU | GAU | CUU |
|  | CheACRV1 | AAUUAUUUU | GAU | CUU |
|  | ChrACRV1 | AAAUAUUUU | GAU | CUU |
|  | ConACRV1 | AAUUCUUUU | GAU | CNC |
|  | CynACRV1 | AAUU(C/A)UUUU | GA(U)_n_ | CNN |
|  | EupACRV1 | AAUUAUUUU | GAU | CUU |
|  | FagACRV1 | AAUUAUUUU | GAU | CNN |
|  | FicACRV1 | AAUUAUUUU | GAU | CNN |
|  | GarACRV1 | AAUUCUUUU | GN(U)_n_ | CNN |
|  | GeuACRV1 | AAUUCUUUU | GAU | CNC |
|  | HedACRV1 | AAUUCUUUU | GNU | CNC |
|  | IleACRV1 | AAUUAUUUU | GA(U)_n_ | CUG |
|  | MedACRV1 | AAUUAUUUU | GAU | CNN |
|  | MenACRV1 | AAUUAUUUU | GAU | CUU |
|  | MorACRV1 | AAUUCUUUU | GNU | CNN |
|  | OakACRV1 | AAUUAUUUU | GAU | CUU |
|  | OciACRV1 | AAUUAUUUU | GAU | CUU |
|  | PelACRV1 | AAUUAUUUU | GAU | CUN |
|  | PhyACRV1 | AAUUCUUUU | GAU | CUC |
|  | PinACRV1 | AAUUAUUUU | GN(U)_n_ | CU(U/G) |
|  | PogACRV1 | AAUUCUUUU | G(N)_n_ | CUC |
|  | PogACRV2 | AAUUAUUUU | GAU | CNN |
|  | PogACRV3 | AAUUAUUUU | GAU | CNG |
|  | PriACRV1 | AAUUCUUUU | GA(U)_n_ | CUN |
|  | PriACRV2 | CAUUAUUUU | GAU | CUG |
|  | RosACRV1 | AAUUAUUUU | GAU | CUN |
|  | RubACRV1 | AAUUAUUUU | GNU | CNN |
|  | ScuACRV1 | AAUUAUUUU | G(N)_n_ | CNN |
|  | TolACRV1 | AAUUCUUUU | GNU | CUC |
|  | TriACRV1 | AAUUAUUUU | GA(U)_n_ | CU(G/U) |
|  | UtrACRV1 | AAUUAUUUU | GA(U)_n_ | CNN |
|  | WMaACRV1 | AAUUCUUUU | GAU | CUU |
|  | WurACRV1 | AAUUAUUUU | GN(U)_n_ | CNN |
|  | ZeaACRV1 | AUUUAUUUU | GA(U)_n_ | CNN |
|  | AcCV | AAUUAUUUU | GAU | CUG |
|  | ADV | AAUUAUUUU | GAU | CUU |
|  | AscSyV1 | AAUUAUUUU | GNU | CNN |
|  | BCRV2 | AAUUCUUUU | GNU | CNN |
|  | BmV1 | AAUUAUUUU | GAN | CUG |
|  | CCyV1 | AAUUCUUUU | G(N)_n_ | CUU |
|  | ChYDaV | AAUUAUUUU | GAU | CUN |
|  | CCRV1 | AAUUAUUUU | GAU | CUU |
|  | CnV2 | AAUUAUUUU | GAU | CUN |
|  | DV1 | AAUUAUUUU | GAU | CUG |
|  | GlLV1 | AAUUAUUUU | GAU | CUU |
|  | HpLV | AAUUAUUUU | GAU | CNN |
|  | KePCyV | AAUUAUUUU | GAU | CUU |
|  | LNYV | AAUUCUUUU | G(N)_n_ | CUU |
|  | LYMoV | AAUUCUUUU | G(N)_n_ | CUN |
|  | NymAV1 | AUUAAUUUU | GAU | CUN |
|  | PaCRV1 | AAUUAUUUU | GAU | CUU |
|  | PCaCV | AAUUAUUUU | GNU | CUN |
|  | PeVA | AAUUAUUUU | G(N)_n_ | CUN |
|  | PNSaV | AAUUAUUUU | GAU | CUN |
|  | RVCV | AUUUAUUUU | GAU | CUU |
|  | SaV1 | AUUUAUUUU | GAU | CNN |
|  | SCV | AAUUAUUUU | GAU | CUU |
|  | StrV1 | AAUUAUUUU | GAU | CUU |
|  | StrV2 | AAUUCUUUU | GNU | CNN |
|  | TCRV1 | AAUUAUUUU | GAU | CNN |
|  | TpVA | AAUUAUUUU | GAU | CUU |
|  | TpVB | AAUUCUUUU | G(N)_n_ | CUN |
|  | TrARV1 | AAUUAUUUU | GAU | CUU |
|  | TYMaV | AAUUAUUUU | GAU | CUU |
|  | WhIV4 | AAUUAUUUU | GNU | CUU |
|  | WhIV5 | AAUUAUUUU | GAU | CNN |
|  | WhIV6 | AAUUAUUUU | GAU | CUN |
| *Betacytorhabdovirus* | ArtBCRV1 | AUUCUUUUU | GUU | CUU |
|  | BegBCRV1 | AUAUUUUUU | GN | CUN |
|  | BetBCRV1 | AUUCUUUUU | GG(U)_n_ | CUG |
|  | BetBCRV2 | AUUCUUUUU | GG(U)_n_ | CUG/A |
|  | BouBCRV1 | AUUCUUUUU | GCU | CUG |
|  | ChrBCRV1 | AUUCUUUUU | GUU | CUU |
|  | CorBCRV1 | AUUCUUUUU | GGUU | CUG |
|  | CucBCRV1 | AUUCUUUUU | G(N)_n_ | CUU |
|  | CypBCRV1 | UUCUUUUUU | GA | CUC |
|  | DryBCRV1 | AUUAUUUUU | GGU | CCU |
|  | DurBCRV1 | AUUCUUUUU | GA | CUC |
|  | GleBCRV1 | AUUAUUUUU | GG(U)_n_ | CUN |
|  | GlyBCRV1 | AUUAUUUUU | GGU | CCU |
|  | HepBCRV1 | AUUAUUUUU | GA(U)_n_ | CUU |
|  | HowBCRV1 | AUAUUUUUU | GA | CUN |
|  | IpoBCRV1 | AUUCUUUUU | GUU | CUN |
|  | JusBCRV1 | AUU(A/C)UUUUU | GGUU | CUN |
|  | KobBCRV1 | AUUCUUUUU | GGN | CUC |
|  | Leu CRV1 | AUUCUUUUU | GA | CUC |
|  | LycBCRV1 | AUUAUUUUU | GGU | CCU |
|  | ManBCRV1 | AUUAUUUUU | GG(U)_n_ | CUN |
|  | MorBCRV1 | AUUAUUUUU | GGU | CCU |
|  | NitBCRV1 | AUUCUUUUU | GGUU | CUN |
|  | PanBCRV1 | AUUCUUUUU | G(G/A) | CUC |
|  | PasBCRV1 | AUAUUUUUU | GAUU | CUC |
|  | PSaBCRV1 | AUUUAUUUU | GA | CUC |
|  | PSaBCRV2 | AUUAUUUUU | GNU | CUN |
|  | PenBCRV1 | AUAUUUUUU | G(N)_n_ | CUU |
|  | PheBCRV1 | AUUAUUUUU | GGUU | CUC |
|  | PopBCRV1 | AUUCUUUUU | GG(U)_n_ | CUN |
|  | PueBCRV1 | AUUAUUUUU | GGU | CCU |
|  | SesBCRV1 | UUCUUUUUU | GA | CUN |
|  | SchBCRV1 | AUUCUUUUU | GA | CUC |
|  | SopBCRV1 | AUUAUUUUU | GGU | CCU |
|  | TriBCRV1 | AUUCUUUUU | GN | CUN |
|  | VicBCRV1 | AUUCUUUUU | GG | CUC |
|  | ZanBCRV1 | AUUAUUUUU | GGU | CCU |
|  | ZanBCRV2 | AUUAUUUUU | GGU | CCU |
|  | ZanBCRV3 | AUUAUUUUU | GGU | CCU |
|  | AntAmV1 | AUUAUUUUU | GCU | CUU |
|  | AriACRV | UUAUUUUUU | GN(N)_n_ | CNN |
|  | BeTaV1 | UUAUUUUUU | GA | CUC |
|  | BYSMV | AUUAUUUUU | GA | CUC |
|  | CBDaV | AUUCUUUUU | GG | CUC |
|  | CuCV1 | AUUAUUUUU | GA | CUC |
|  | MaCyV | AUUCUUUUU | GA | CUC |
|  | MYSV | AUUAUUUUU | GA | CUC |
|  | NCMV | AUUCUUUUU | GA | CUC |
|  | PMuMaV | AUUAUUUUU | G(N)_n_ | CUA |
|  | PpVE | AUUCUUUUU | GAC | CCU |
|  | RaCV | AUUCUUUUU | G(N)_n_ | CUN |
|  | RVR | AUUUAUUUU | GA | CUC |
|  | RSMV | AUUCUUUUU | GCU | CUG |
|  | RudV1 | AUUCUUUUU | GGUU(N)_n_ | CUN |
|  | SbBMV | UUAUUUUUU | GA | CAC |
|  | TaEV1 | AUUCUUUUU | GG(N)_n_ | CUN |
|  | TiCRV1 | AUAUUUUUU | GA(N)_n_ | CUC |
|  | YmCaV | UUAUUUUUU | GA | CUC |
|  | YmVA | AUUCUUUUU | GGU | CCU |
| *Gammacytorhabdovirus* | ArgGCRV1 | AUUCUUUUU | AAU | CCU |
|  | CarGCRV1 | AUUCUUUUU | A(N)_n_ | CCU |
|  | CelGCRV1 | AUUCUUUUU | A(N)_n_ | CNU |
|  | CopGCRV1 | AUUCUUUUU | A(N)_n_ | CCU |
|  | CusGCRV1 | AUUCUUUUU | A(N)_n_ | CNN |
|  | CusGCRV2 | AUUCUUUUU | A(N)_n_ | CCU |
|  | CypGCRV1 | AAUCUUUUU | A(N)_n_ | CNN |
|  | EpiGCRV1 | AUUCUUUUU | AUGU | CCU |
|  | FraGCRV1 | AUUCUUUUU | A(N)_n_ | CNU |
|  | FraGCRV2 | AUUCUUUUU | A(N)_n_ | CCU |
|  | HelGCRV1 | AUUCUUUUU | A(N)_n_ | CCU |
|  | HibGCRV1 | AUUCUUUUU | A(N)_n_ | CNN |
|  | LonGCRV1 | AUUCUUUUU | A(N)_n_ | CCU |
|  | LupGCRV1 | AUUCUUUUU | A(N)_n_ | CCU |
|  | Rh GCRV1 | AUUUCUUUU | A(N)_n_ | CCU |
|  | SilGCRV1 | AUUCUUUUU | A(N)_n_ | CCU |
|  | GymDenV1 | AAUCUUUUU | A(N)_n_ | CNN |
|  | TrAV1 | AUUCUUUUU | A(N)_n_ | CNU |
| *Trirhavirus* | AlTRV1 | AAUUCUUUU | GN(N)_n_ | CUC |
|  | ChTRV1 | AAUUCUUUU | GN(N)_n_ | CCU |
|  | EryTRV1 | AAUUCUUUU | GN(N)_n_ | CUC |
|  | MeTRV1 | AAUUCUUUU | GN(N)_n_ | CU (C/G) |
|  | PiTRV1 | AAUUCUUUU | GN(N)_n_ | CUN |

The consensus gene junction sequences of the viruses identified in this study are highlighted in light grey. * Names and abbreviations of newly identified viruses are listed in Tables 1-4; while the names and abbreviations of known viruses are listed in Supp Table 1.
