## Supplementary Table S1 for "Novel tri-segmented rhabdoviruses: A data mining expedition unveils the cryptic diversity of cytorhabdoviruses"

Supplementary Table S1. Virus names and abbreviations of cytorhabdovirus sequences used in this study.

| **Virus name** | **Abbreviation** |
| --- | --- |
| Actinidia cytorhabdovirus | AcCV |
| alfalfa dwarf virus | ADV |
| Anthurium Amnicola virus 1 | AntAmV1 |
| Aristolochia associated cytorhabdovirus | AriACRV |
| Asclepias syriaca virus 1 | AscSyV1 |
| Bacopa monnieri virus 1 | BmV1 |
| Bemisia tabaci associated virus 1 | BeTaV1 |
| blackcurrant cytorhabdovirus 2 | BCRV2 |
| barley yellow striate mosaic virus | BYSMV |
| cabbage cytorhabdovirus 1 | CCyV1 |
| chrysanthemum yellow dwarf-associated virus | ChYDaV |
| Cirsium cytorhabdovirus 1 | CCRV1 |
| Cnidium virus 2 | CnV2 |
| Colocasia bobone disease-associated virus | CBDaV |
| cucurbit cytorhabdovirus 1 | CuCV1 |
| Daphne virus 1 | DV1 |
| Glehnia littoralis virus 1 | GlLV1 |
| Gymnadenia densiflora virus 1 | GymDenV1 |
| Hyptis latent virus | HpLV |
| Kenyan potato cytorhabdovirus | KePCyV |
| lettuce necrotic yellows virus | LNYV |
| lettuce yellow mottle virus | LYMoV |
| maize associated cytorhabdovirus | MaCyV |
| maize yellow striate virus | MYSV |
| northern cereal mosaic virus | NCMV |
| Nymphaea alba virus 1 | NymAV1 |
| papaya virus E | PpVE |
| paper mulberry mosaic-associated virus | PMuMaV |
| Pastinaca cytorhabdovirus 1 | PaCRV1 |
| patchouly chlorosis associated virus | PCaCV |
| persimmon virus A | PeVA |
| Physostegia chlorotic mottle virus | PhCMoV |
| Plumbago necrotic spot associated virus | PNSaV |
| potato yellow dwarf virus | PYDV |
| raspberry vein chlorosis virus | RVCV |
| rice stripe mosaic virus | RSMV |
| rose associated cytorhabdovirus | RaCV |
| rose virus R | RVR |
| Rudbeckia virus 1 | RudV1 |
| Sambucus virus 1 | SaV1 |
| strawberry crinkle virus | SCV |
| strawberry virus 1 | StrV1 |
| strawberry virus 2 | StrV2 |
| soybean blotchy mosaic virus | SbBMV |
| Tagetes erecta virus 1 | TaEV1 |
| Taraxacum cytorhabdovirus | TCRV1 |
| Tillia cytorhabdovirus 1 | TiCRV1 |
| tomato yellow mottle associated virus | TYMaV |
| Trachyspermum ammi virus 1 | TrAV1 |
| Trichosanthes-associated rhabdovirus 1 | TrARV1 |
| Trifolium pratense virus A | TpVA |
| Trifolium pratense virus B | TpVB |
| Wuhan insect virus 4 | WhIV4 |
| Wuhan insect virus 5 | WhIV5 |
| Wuhan insect virus 6 | WhIV6 |
| yerba mate chlorosis-associated virus | YmCaV |
| yerba mate virus A | YmVA |
