## Supplementary figures and images for "Novel tri-segmented rhabdoviruses: A data mining expedition unveils the cryptic diversity of cytorhabdoviruses"

### Fig. S1

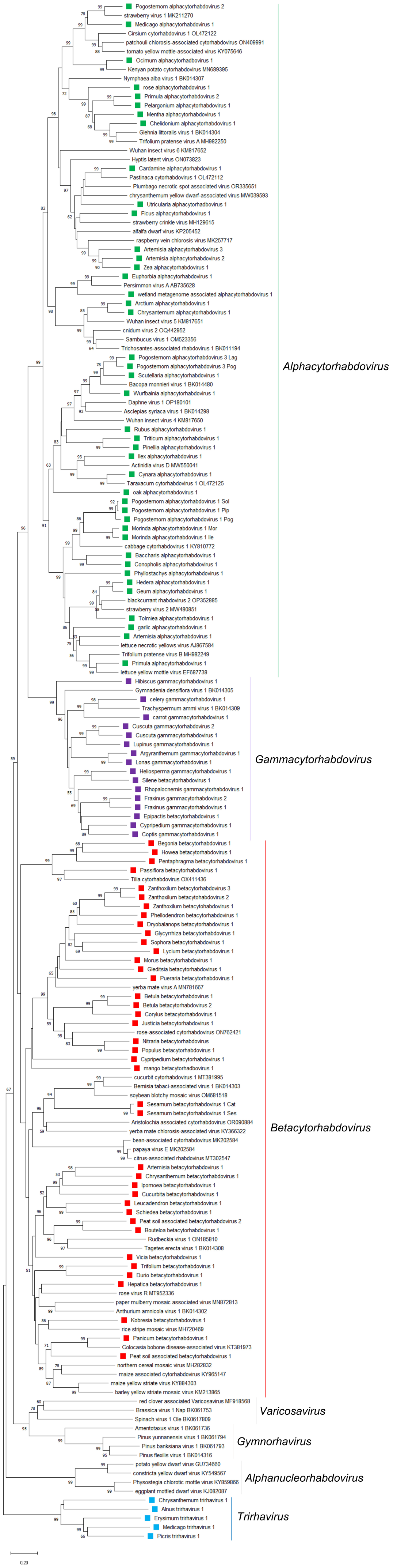
